# Intrinsic excitability in layer IV-VI anterior insula to basolateral amygdala projection neurons encodes the confidence of taste valence

**DOI:** 10.1101/2022.05.23.493046

**Authors:** Sailendrakumar Kolatt Chandran, Adonis Yiannakas, Haneen Kayyal, Randa Salalha, Federica Cruciani, Liron Mizrahi, Mohammad Khamaisy, Shani Stern, Kobi Rosenblum

## Abstract

Avoiding potentially harmful, and consuming safe food is crucial for the survival of living organisms. However, sensory information can change its valence following conflicting experiences. Novelty and aversiveness are the two crucial parameters defining the currently perceived valence of taste. Importantly, the ability of a given taste to serve as CS in conditioned taste aversion (CTA) is dependent on its valence. Activity in anterior insula (aIC) layer IV-VI pyramidal neurons projecting to the basolateral amygdala (BLA) is correlative and necessary for CTA learning and retrieval, as well as the expression of neophobia towards novel tastants, but not learning taste familiarity. Yet, the cellular mechanisms underlying the updating of taste valence representation in this specific pathway are poorly understood. Here, using retrograde viral tracing and whole cell patch-clamp electrophysiology in trained mice, we demonstrate that the intrinsic properties of deep-lying layer IV-VI, but not superficial layer I-III aIC-BLA neurons, are differentially modulated by both novelty and valence, reflecting the subjective predictability of taste valence arising from prior experience. These correlative changes in the profile of intrinsic properties of LIV-VI aIC-BLA neurons were detectable following both simple taste experiences, as well as following memory retrieval, extinction learning and reinstatement.

## Introduction

In the natural setting, animals approach novel taste stimuli tentatively, as to closely examine them according to a genetic plan, as well as in relation to associated visceral consequences (Schier and Spector, 2019). Bitter and sour tastes are innately aversive, acting as warning signals for the presence of toxins (Bachmanov et al., 1996). Conversely, neophobia to innately appetitive sweet and moderately salty tastants dissipates over time (Lin et al., 2012). Importantly, animals can learn to avoid innately appetitive tastants (e.g., saccharin-, or NaCl-water – the conditioned stimulus, CS), through conditioned taste aversion - CTA (Garcia et al., 1955; Nachman and Ashe, 1973). This single-trial associative learning paradigm results in robust aversion following the pairing of the CS with a malaise-inducing agent (the unconditioned stimulus, US), such as LiCl (Bures et al., 1998). CTA memories are robust, but can be extinguished through unreinforced CS re-exposures, and subsequently reinstated through US re-exposure (Schachtman et al., 1985; Mickley et al., 2004). Unlike other forms of classical conditioning, the inter-stimulus interval (ISI) between taste experience (CS) and visceral outcome (US), extends to several hours (Adaikkan and Rosenblum, 2015). How CTA learning enables this long-trace associative process, within timeframes that deviate from classical Hebbian plasticity mechanisms is currently unknown (Chinnakkaruppan et al., 2014; Adaikkan and Rosenblum, 2015).

The primary taste cortex - the anterior insula (aIC), along with the basolateral amygdala (BLA), govern the encoding and retrieval of taste information (Piette et al., 2012; Bales et al., 2015). Gustatory processing at the IC encompasses thalamocortical and corticocortical inputs that relay taste-, as well as palatability-related inputs from the BLA, that reflect the emotional valence associated with taste stimuli (Stone et al., 2020). Neuronal taste responses at the IC and BLA are using temporal information to encode multiple types of information relating to stimulus identity and palatability (Grossman et al., 2008; Sadacca et al., 2012; Arieli et al., 2020; Vincis et al., 2020). Both synaptic plasticity and neuronal intrinsic properties are proposed to serve as cellular mechanisms underlying learning and memory (Citri and Malenka, 2008; Sehgal et al., 2013). CTA learning promotes LTP induction in the BLA-IC pathway (Jones et al., 1999; Juárez-Muñoz et al., 2017), and strengthens cell-type specific functional connectivity along the projection (Haley et al., 2016). Intrinsic excitability is the tendency of neurons to fire action potentials when exposed to inputs, reflecting changes in the suit and properties of specific ion channels (Disterhoft et al., 2004; Song and Moyer, 2018). Even though independent mechanisms are involved, recent evidence indicates learning and memory necessitates the that coupling of intrinsic and synaptic plasticity (Turrigiano, 2011; Greenhill et al., 2015; Wu et al., 2021).

The IC is an integration hub tuned for the encoding of both exteroceptive as interoceptive information (Gogolla et al., 2014; Haley and Maffei, 2018; Livneh et al., 2020; Koren et al., 2021). By virtue of its extensive network of connectivity, this elongated cortical structure has been shown to integrate sensory, emotional, motivational, and cognitive brain centers through distinct mechanisms. For example, deletions of either *Fos* or *Stk11* in BLA-aIC neurons, alter intrinsic properties at the aIC, and impair CTA acquisition (Levitan et al., 2020). Furthermore, approach behaviors in social decision-making are modulated by subjective and sex-specific affective states that regulate cell-type-specific changes in intrinsic properties at IC terminals receiving hypothalamic input (Rogers-Carter et al., 2018, 2019; Rieger et al., 2022). The posterior IC (pIC) integrates visceral-sensory signals of current physiological states with hypothalamus-gated amygdala anticipatory inputs relating to food or water ingestion, to predict future physiological states (Livneh et al., 2017, 2020). Conversely, aversive visceral stimuli such as LiCl, activate CaMKII neurons projecting to the lateral hypothalamus in right-, but not the left IC, whose optogenetic activation or inhibition can bidirectionally regulate food consumption (Wu et al., 2020). We have previously shown that the aIC-BLA projection is necessary and sufficient for CTA acquisition and retrieval (Lavi et al., 2018; Kayyal et al., 2019), while CTA retrieval requires activation of the projection concomitant with parvalbumin (PV) interneurons (Yiannakas et al., 2021). Moreover, artificial activation of aIC-BLA projecting neurons is sufficient to induce CTA for appetitive taste (Kayyal et al., 2019). Here, using retrograde viral tracing, behavioral analysis, and whole-cell patch-clamp slice electrophysiology, we assessed two hypotheses: (1) That the intrinsic properties of the aIC-BLA projection change as a function of certainty of taste valence prediction, but not percept; and (2) that predictive valence-specific changes in intrinsic properties would be reflected through excitability, being low when taste outcome is highly predictive (i.e., following CTA retrieval or unreinforced familiarization), and high when taste valence is uncertain (i.e., following novelty or extinction). Our data demonstrate for the first time that the intrinsic properties of LIV-VI aIC-BLA neurons are differentially regulated by innate and learned drives, reflecting the confidence of currently perceived taste valence.

## Materials and methods

### Animals

Animals used were 8–12-week-old C57BL/6j (WT) adult male mice. Mice were kept in the local animal resource unit at the University of Haifa on a 12-hour dark/light cycle. Water and chow pellets were available ad libitum, while ambient temperature was tightly regulated. All procedures conducted were approved by the University of Haifa Animal Care and Use Committee (Ethics License 554/18), as prescribed by the Israeli National Law for the Protection of Animals – Experiments with Animals (1994).

### Animal surgery and viral injections

Following surgery and stereotactic injection of viral vectors, behavioral paradigms were performed, as previously described (Yiannakas et al., 2021). Briefly, mice were treated with norocarp (0.5mg/kg), before being anesthetized (M3000 NBT Israel/Scivena Scientific) and transferred to a Model 963 Kopf® stereotactic device. Upon confirming the lack of pain responses, the skull was surgically exposed and drilled to bilaterally inject 0.25μl of ssAAV_retro2-hSyn1-chi-mCherry-WPRE-SV40p(A) (physical titer 8.7 x 10E12 vg/ml), at the BLA (AP -1.58; ML ± 3.375; DV - 4.80). Viral delivery was performed using a Hamilton micro-syringe (0.1uL/minute), while the sculp was cleaned and closed using Vetbond®. Animals were then administered with 0.5mg/kg norocarp and 0.5mg/kg of Baytril (enrofloxacin), and then transferred to a clean and heat-adjusted enclosure for 2 hours. Upon inspection, mice were returned to fresh cages along with similarly treated cage-mates. Weight-adjusted doses of the Norocarp and Baytril were administered for an additional 3 days. All AAV constructs used in this study were obtained from the Viral Vector Facility of the University of Zurich (http://www.vvf.uzh.ch/).

### Electrophysiological studies of the influence of innate taste identity, novelty, and valence on aIC-BLA excitability

WT mice treated with viral constructs labeling aIC-BLA projecting neurons were used for electrophysiological studies. Upon recovery, mice were randomly assigned into treatment groups (Figure 1). Following 24hrs of water deprivation, animals were water restricted for 3 days, receiving water in pipettes ad libitum for 20 minutes/day (Kayyal et al., 2019; Yiannakas et al., 2021). This regime has been extensively used by our lab as it allows rodents to reliably learn to drink from water pipettes with minimal weight loss. Mean total drinking was recorded on the 3^rd^ day of water restriction. Novel taste consumption groups were presented with 1.0mL of either 0.5% saccharin (*Saccharin 1x*), or Quinine 0.014% (*Quinine 1x*). One hour following the final taste presentation, animals were subjected to patch-clamp electrophysiology (Kayyal et al., 2021; Yiannakas et al., 2021). The *Water* group underwent the same behavioral procedure without novel taste presentations were sacrificed for electrophysiological investigations one hour following water presentation. To dissociate between taste identity and familiarity-related changes in electrophysiological properties, a cohort of mice treated to label the aIC-BLA projection were similarly water deprived following familiarization with saccharin (*Saccharin 5x*). Following the initial water restriction, *Saccharin 5x* animals were allowed access to 0.5% saccharin, in 20minute sessions for 4 days. On the fifth day, mice were provided with 1.0ml of the tastant, 1 hour prior to sacrifice for electrophysiological recordings. Additionally, WT animals injected with the same viral vector, were allowed a month to recover, following which they were sacrificed for electrophysiological investigations without any behavioral manipulation (*Cage Controls*).

**Figure 1:**
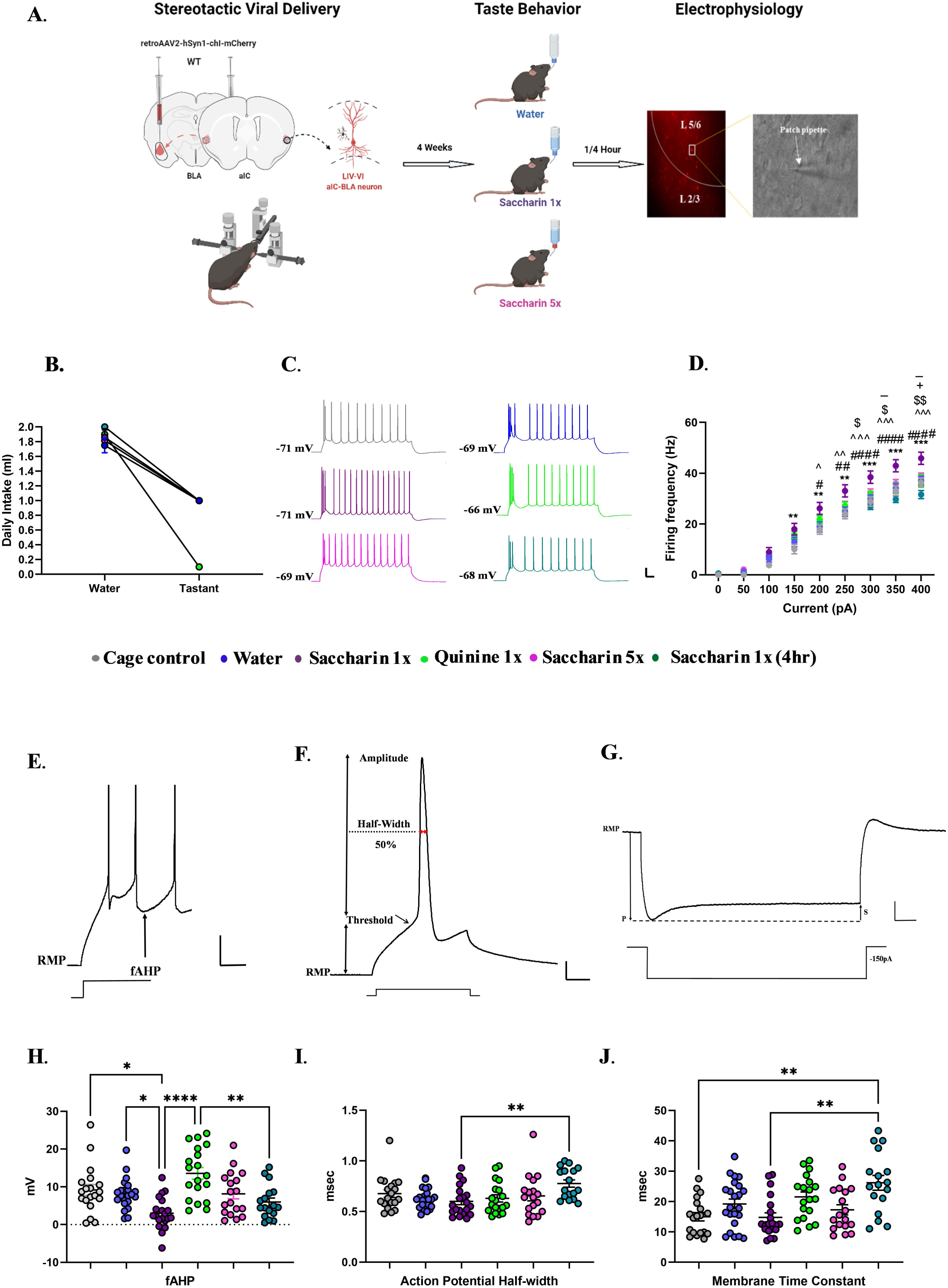
Retrieval of appetitive and novel taste increases excitability in LIV-VI aIC-BLA projection neurons A) Diagrammatic representation of experimental procedures. Following surgery and stereotaxic delivery of ssAAV_retro2-hSyn1-chi-mCherry-WPRE-SV40p(A) into the BLA, mice were allowed 4 weeks of recovery. Animals were subsequently assigned to treatment groups and trained to drink from pipettes (see Methods). We compared the intrinsic properties of LIV-VI aIC-BLA neurons among the Water (n=6 animals, 23 cells), Saccharin 1x (n=5 animals, 20 cells), Saccharin 1x(4h) (n=4 animals, 17 cells), Saccharin 5x (n=6 animals, 18 cells) and Quinine 1x groups (n=4 animals, 19 cells), as well as a Cage control group (n=4 animals, 19 cells) that underwent surgery and stereotaxic delivery of ssAAV_retro2-hSyn1-chi-mCherry-WPRE-SV40p(A) at the BLA without water restriction. B) Graph showing the water consumption prior to treatment (mean ± SD). There was no significant difference between water intakes between the groups before the treatment. One-Way ANOVA, p= 0.9766. C) Representative traces of LIV-VI aIC-BLA projecting neurons from the six treatment groups. Scale bars 20 mV vertical and 50ms horizontal from 300 pA step. D) The dependence of firing rate on current step magnitude in LIV-VI aIC-BLA neurons was significantly different among the treatment groups. Excitability in the Saccharin 1x was increased compared to all other groups. Two-way repeated measures ANOVA, Current x Treatment: p<0.0001; Cage control vs. Saccharin 1x: **p<0.01, ***p<0.001; Saccharin 1x vs. Saccharin 1x (4hr) : #p<0.05, ##p<0.01, ####p<0.0001; Water vs. Saccharin 1x: ^p<0.05, ^^p<0.01, ^^^ p<0.001; Saccharin 1x vs. Quinine 1x: $ p<0.05, $$p<0.01; Saccharin 1x vs. Saccharin 5x: - p<0.05; Saccharin 1x (4hr) vs. Saccharin 5x : + p<0.05. E) Representative of all fAHP measurements in response to 500 msec step current injections. Scale bars 20 mV vertical and 50 msec horizontal. F) Representative of all action potential properties were taken. Scale bars 20 mV vertical and 5 msec horizontal. G) Measurements for all input resistance, sag ratio and membrane time constants were analyzed in response to 1 sec, -150pA step current injection. P- peak voltage, S- steady state voltage. Scale bars 5 mV vertical and 100 msec horizontal. H) Significant differences were observed among the treatment groups in terms of fAHP. Cage control (9.191 ± 1.449 mV), Water (8.150 ± 0.8288 mV), Saccharin 1x (3.016 ± 0.9423 mV), Quinine 1x (13.58 ± 1.562 mV) Saccharin 5x (8.158 ± 1.356 mV), Saccharin 1x (4hrs) (5.989 ± 1.074 mV), One-Way ANOVA, p<0.0001. I) Action potential half-width in the Saccharin 1x group (0.6005 ± 0.03260 msec) was significantly decreased compared to Saccharin 1x (4hr) 90.7765 ± 0.03641 msec), One-Way ANOVA, p=0.0065. J) The membrane time constant was significantly different between the Saccharin 1x (14.82 ± 1.485 msec) and Saccharin 1x (4hrs) (26.21 ± 2.421 msec) groups and Cage control (15.03 ± 1.376 msec), One-Way ANOVA, p=0.0005. For panels 1D, H, I and J: *p<0.05, **p<0.01, ***p<0.001, ****p<0.0001. All data are shown as mean ± SEM.

### Electrophysiological studies of the influence of learned aversive taste memory retrieval on aIC-BLA excitability

WT mice were treated with viral constructs labelling aIC-BLA projecting neurons to assess the electrophysiological properties of the projection during aversive or appetitive taste memory retrieval. Upon recovery, mice in CTA retrieval group were trained in CTA for saccharin (LiCl 0.14M, 1.5% body weight), while the appetitive saccharin retrieval group (*Saccharin 2x*) received a matching body weight adjusted injection of saline (Yiannakas et al., 2021). Three days following conditioning, both groups underwent a memory retrieval task, receiving 1.0mL of the conditioned tastant 1 hour prior to sacrifice (Figures 2, 4). Brain tissue was extracted and prepared for electrophysiological recording, as above.

**Figure 2:**
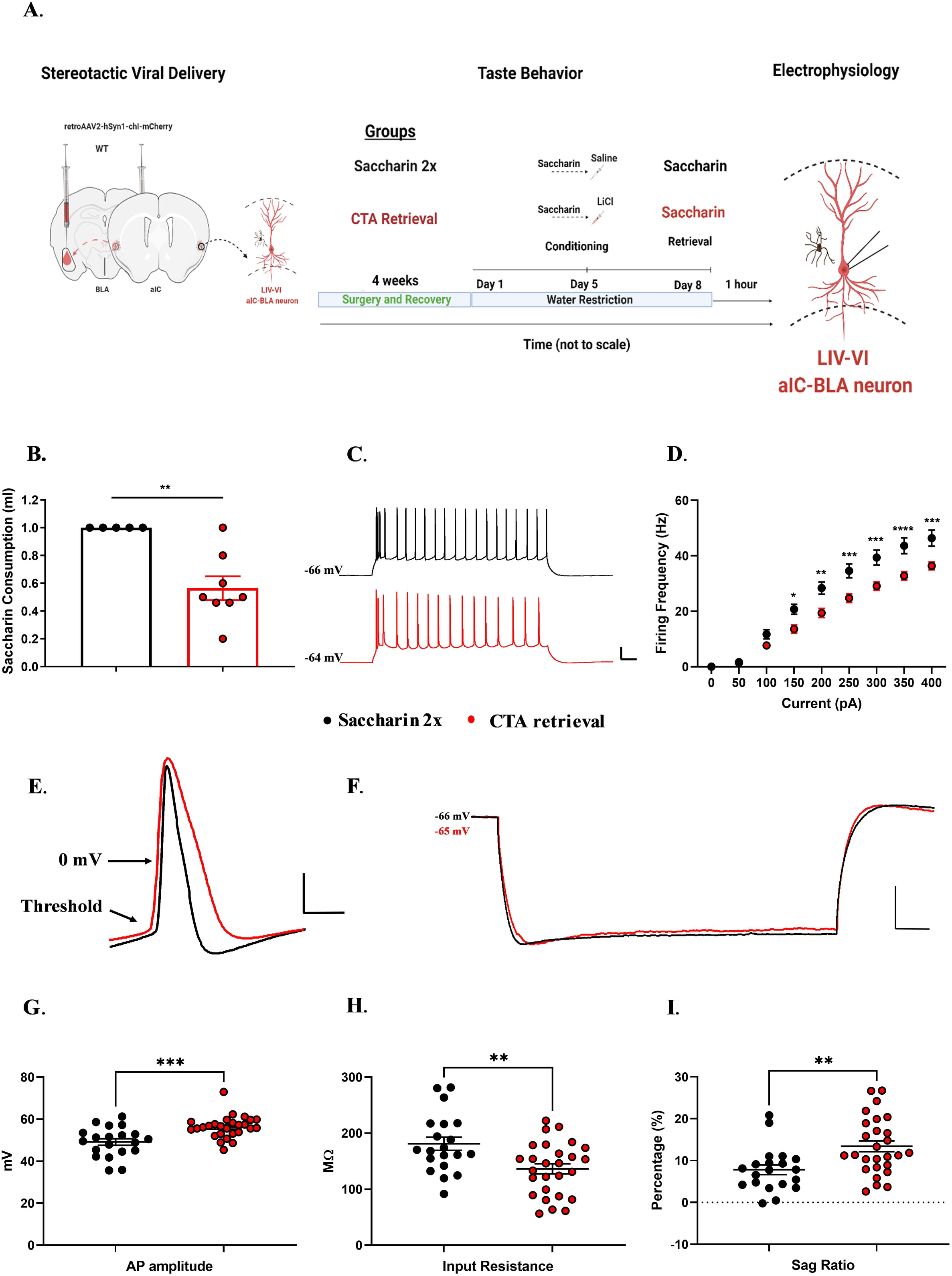
Learned aversive taste memory retrieval suppresses the excitability of LIV-VI aIC-BLA projecting neurons A) Experimental design of behavioral procedures conducted to compare the intrinsic properties of LIV-VI aIC-BLA neurons following learned aversive taste memory retrieval (CTA retrieval - n=8 animals, 27 cells), appetitive retrieval for the same tastant (Saccharin 2x - n=5 animals, 20 cells). B) Mice showed a significantly reduced saccharin consumption following learned aversive memory retrieval (N=8) compared to appetitive retrieval mice (n=5) group. p=0.0085, Mann Whitney test. C) Representative traces of LIV-VI aIC-BLA projecting neurons from the two treatment groups. Scale bars 20 mV vertical and 50ms horizontal from 300 pA step. D) The excitability of LIV-VI aIC-BLA in the Saccharin 2x group was significantly enhanced compared to CTA retrieval. Two-way repeated measures ANOVA, Current x Treatment: p<0.0001. E) Representative traces showing action potential measurements for both groups. Scale bar 20 mV vertical and 2ms horizontal. F) Representative traces showing the input resistance and sag ratio measurements. Scale bar 10 mV vertical and 100ms horizontal. G) Action potential amplitude in the CTA retrieval (56.21 ± 0.9978 mV) group was increased compared to Saccharin 2x (49.14 ± 1.568 mV), p=0.0005, Mann Whitney test. H) Input resistance in the CTA retrieval group (136.4 ± 9.064 MΩ) was significantly decreased compared to Saccharin 2x (181.1 ± 11.7 MΩ). p=0.0036, Unpaired t test. I) SAG ratio following CTA retrieval (13.41 ± 1.31) was significantly enhanced compared to Saccharin 2x (7.815 ± 1.176). p=0.0037, Unpaired t test. Data are shown as mean ± SEM. *p<0.05, **p<0.01, ***p<0.001, ****p<0.0001.

### Electrophysiological studies of the influence of learned aversive taste memory extinction and reinstatement on aIC-BLA excitability

Electrophysiological studies of CTA extinction and reinstatement were conducted in a cohort of WT male mice (Yiannakas et al., 2021). Following surgery, recovery and water restriction, animals were randomly assigned to the extinction and reinstatement groups (Figures 3-4). Adult male mice used to study extinction and reinstatement were trained in CTA for saccharin following extinction, the reinstatement group received an identical intraperitoneal dose to the original unconditioned stimulus (LiCl 0.14M, 1.5% body weight), 24 hours prior to retrieval. Conversely, the extinction group received a similarly weight-adjusted dose of saline. During the final retrieval session, both groups of mice were allowed access to 1.0mL of the CS, 1 hour prior to sacrifice under deep anesthesia and slice preparation for electrophysiology.

**Figure 3:**
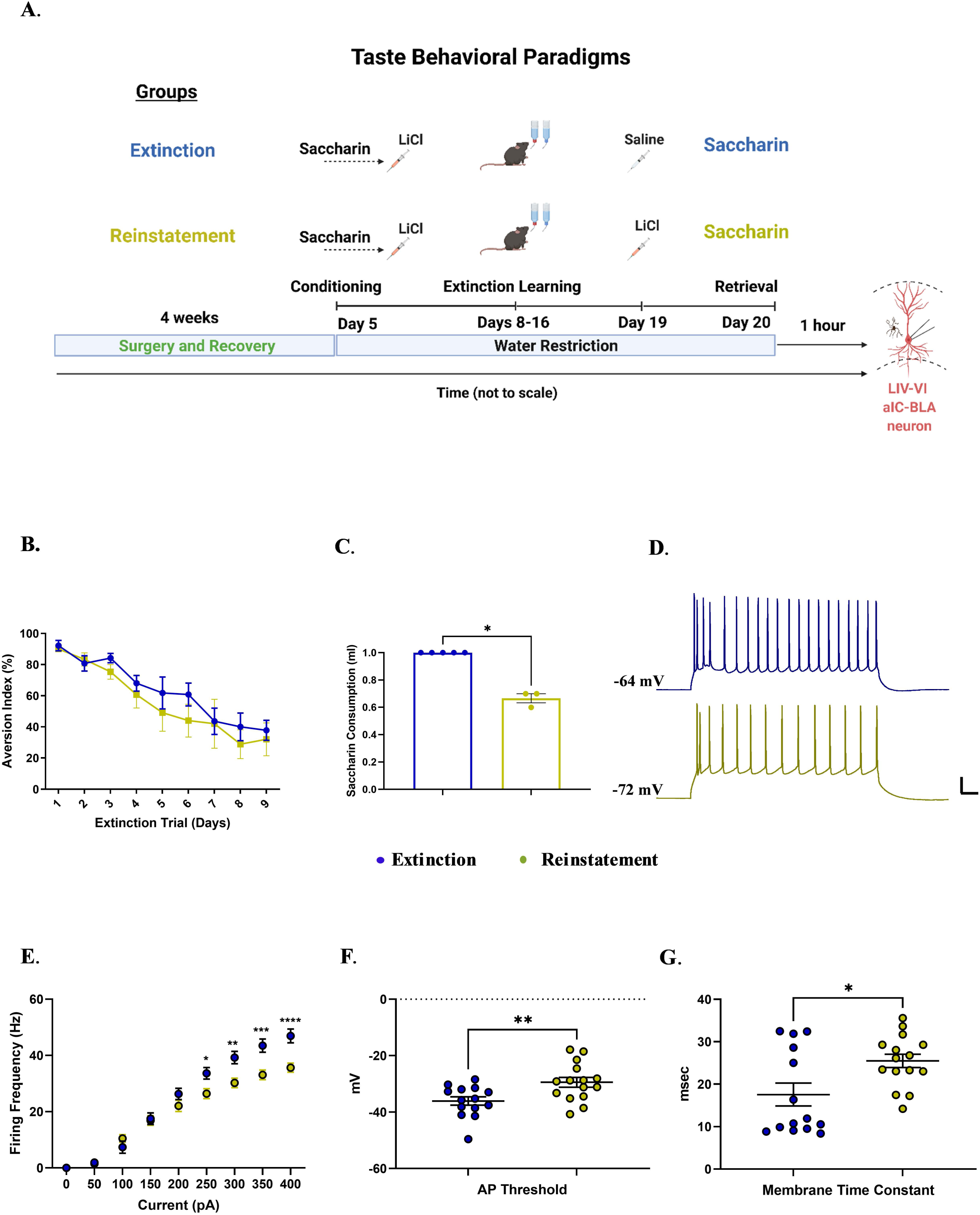
Extinction of CTA enhances, whereas reinstatement suppresses the excitability of LIV-VI aIC-BLA projecting neurons A) Experimental design of behavioral procedures conducted to compare the intrinsic properties of LIV-VI aIC-BLA neurons following CTA Extinction (n=5, animals, 14 cells) and Reinstatement (n=3 animals, 15 cells). B) The graph showing the reduced aversion following the successful extinction in both treatment groups. C) Data showing the saccharin consumption on the test day following successful extinction and Reinstatement of CTA. CTA reinstated mice showed significantly reduced saccharin consumption compared to extinguished mice. p=0.0179, Mann Whitney test. D) Representative traces of LIV-VI aIC-BLA projection neurons firing from the two treatment groups. Scale bars 20 mV and 50ms horizontal from 300 pA step. E) Excitability in LIV-VI aIC-BLA neurons was significantly different among the treatment groups. Two-Way repeated measures ANOVA, Current x Treatment: p<0.0001. F) Action potential threshold in the Reinstatement group (−29.43 ± 1.731 mV) was enhanced compared to Extinction (−36.06 ± 1.481 mV). p=0.0076, Unpaired t test. G) The membrane time constant following Reinstatement (25.48 ± 1.58 msec) was significantly enhanced compared to Extinction (17.55 ± 2.684 msec, p=0.047). p=0.0153, Unpaired t test. For panels 5D-F: *p<0.05, **p<0.01, ***p<0.001, ****p<0.0001. All data are shown as mean ± SEM.

**Figure 4:**
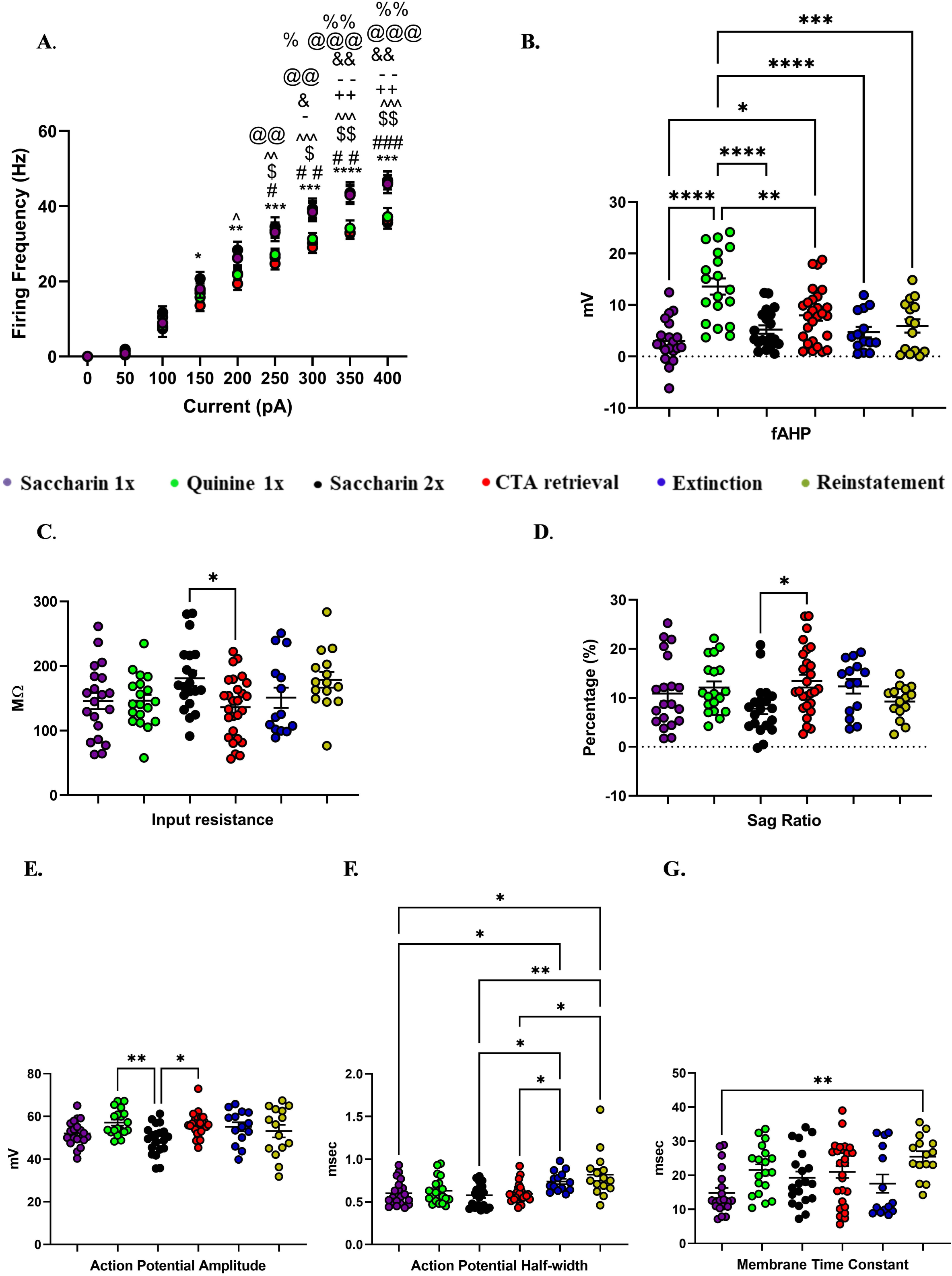
Innately aversive taste is correlated with high fAHP, and prolonged conflicting experiences is correlated with an increased AP half-width in LIV-VI aIC-BLA projecting neurons We compared the intrinsic properties of LIV-VI aIC-BLA neurons among the Saccharin 1x (n=5 animals, 20 cells), Quinine 1x (n=4 animals, 19 cells), Saccharin 2x (n=5 animals, 20 cells), CTA retrieval (n=8, 27 cells), Extinction (n=5 animals, 14 cells) and Reinstatement (n=3 animals, 15 cells) groups. A) Groups associated with positive taste valence (Saccharin 1x, Saccharin 2x, Extinction), exhibited significantly increased excitability compared to innate or learned negative taste valence groups (Quinine 1x, CTA retrieval and Reinstatement). Two-way repeated measures ANOVA, Current x Treatment: p<0.0001; Saccharin 2x vs. CTA retrieval: **\***p; Saccharin 2x vs. Reinstatement: **#**p: Saccharin 2x vs Quinine 1x: p**$**; Saccharin 1x vs. CTA retrieval: p**^**; Saccharin 1x vs. Quinine 1x: p**%**; Saccharin 1x vs reinstatement: p**+**; Extinction vs. CTA retrieval: p**@**; Extinction vs. Reinstatement: p**&**; Extinction vs. Quinine 1x: p**-**. B) fAHP was significantly enhanced in response to Quinine 1x (13.56± 1.562 mV) compared to all other groups. Significant differences were also observed between Saccharin 1x (3.016 ± 0.9423 mV), Saccharin 2x (5.223 ± 0.8217 mV), and CTA retrieval (7.97 ± 1.018 mV, p=0.0036). Extinction (4.731 ± 1.021 mV) and Reinstatement (5.932 ± 1.292 mV). One-Way ANOVA, p<0.0001. C) Input resistance was significantly different between Saccharin 2x (181.1 ± 11.7 MΩ) and CTA retrieval (136.4 ± 9.064 MΩ), p= 0.0352. Conversely, Input resistance in Saccharin 1X (145.8 ± 12.56), Quinine 1X (146 ± 9.094), Extinction (151.1 ± 15.63), and Reinstatement groups was similar. One-Way ANOVA, p= 0.0213. D) SAG ratio was significantly different between Saccharin 2x (7.815 ± 1.176) and CTA retrieval (13.41 ± 1.31), p= 0.0209. Conversely, SAG ratio in Saccharin 1x (10.89 ± 1.621), Quinine 1x (12.13 ± 1.23), Extinction (12.37 ± 1.471) and Reinstatement (9.245 ± 0.884) groups was similar One-Way ANOVA, p= 0.0286. E) Action potential amplitude in the Quinine 1x group (57.11 ± 1.376 mV), and CTA retrieval (56.21 ± 0.9978 mV), was significantly increased compared to Saccharin 2x (49.14 ± 1.568 mV, p= 0.0175, and 0.0229, respectively). Conversely, action potential attitude in the Saccharin 1x (52.03 ± 1.308 mV), Extinction (55.09 ± 2.122 mV) and Reinstatement (53.1 ± 2.906 mV) groups was similar. One-Way ANOVA, p = 0.0061. F) Action potential half-width following Extinction (0.7386 ± 0.03145 msec) and Reinstatement (0.8187 ± 0.06929 msec) was elevated compared to Saccharin 1x (0.6005 ± 0.03260 msec), Saccharin 2x (0.5780 ± 0.02994 msec) as well as CTA retrieval (0.5959 ± 0.02080 msec, but no with Quinine 1x (0.6300 ± 0.03555 msec). One-Way ANOVA, p = 0.0002. G) The membrane time constant in the Saccharin 1x (14.82 ± 1.485 msec) group was significantly suppressed compared to Reinstatement (25.48 ± 1.58 msec, p= 0.0043) groups was. Differences between CTA retrieval (20.96 ± 1.724 msec, p=0.0189), Quinine 1x (21.55 ± 1.638 msec), Saccharin 2x (19.28 ± 1.837 msec) and Extinction (17.55 ± 2.684 msec) groups failed to reach significance. One-Way ANOVA, p = 0.0047.

### Electrophysiology tissue preparation

The slice electrophysiology and recording parameters were used as described previously (Kayyal et al., 2021; Yiannakas et al., 2021). Briefly, mice were deeply anesthetized using isoflurane, while brains were extracted following decapitation. Three-hundred um thick coronal brain slices were obtained with a Campden-1000® Vibratome. Slices were cut in ice-cold sucrose-based cutting solution containing the following (in mM): 110 sucrose, 60 NaCl, 3 KCl, 1.25 NaH2PO4, 28 NaHCO3, 0.5 CaCl2, 7 MgCl2, 5 D-glucose, and 0.6 ascorbate. The slices were allowed to recover for 30 min at 37°C in artificial CSF (ACSF) containing the following (in mM): 125 NaCl, 2.5 KCl, 1.25 NaH2PO4, 25 NaHCO3, 25 D-glucose, 2 CaCl2, and 1 MgCl2. Slices were then kept for an additional 30 min in ACSF at room temperature until electrophysiological recording. The solutions were constantly gassed with carbogen (95% O2, 5% CO2).

### Intracellular whole cell recording

After the recovery period, slices were placed in the recording chamber and maintained at 32-34°C with continuous perfusion of carbogenated ACSF (2 ml/min). Brain slices containing the anterior insular cortices were illuminated with infrared light and pyramidal cells were visualized under a differential interference contrast microscope with 10X or 40X water-immersion objectives mounted on a fixed-stage microscope (BX51-WI; Olympus®). The image was displayed on a video monitor using a charge-coupled device (CCD) camera (QImaging®, Canada). Insula to BLA projection cells infected with AAV were identified by visualizing mCherry^+^ cells. Recordings were amplified by Multiclamp™ Axopatch™ 200B amplifiers and digitized with Digidata® 1440 (Molecular Devices®). The recording electrode was pulled from a borosilicate glass pipette (3–5 M) using an electrode puller (P-1000; Sutter Instruments®) and filled with a K-gluconate-based internal solution containing the following (in mM): 130 K-gluconate, 5 KCl, 10 HEPES, 2.5 MgCl2, 0.6 EGTA, 4 Mg-ATP, 0.4 Na3GTP and 10 phosphocreatine (Na salt). The osmolarity was 290 mOsm, and pH was 7.3. The recording glass pipettes were patched onto the soma region of mCherry^+^ pyramidal neurons and neighboring non fluorescent pyramidal neurons.

The recordings were made from the soma of insula pyramidal cells, particularly from layer 2/3 and Layer 5/6. Liquid junction potential (10 mV) was not corrected online. All current clamp recordings were low pass filtered at 10 kHz and sampled at 50 kHz. Pipette capacitance and series resistance were compensated and only cells with series resistance smaller than 20 MΩ were included in the dataset. The method for measuring active intrinsic properties was based on a modified version of previous protocols (Kaphzan et al., 2013; Chakraborty et al., 2017; Sharma et al., 2018).

### Recording parameters

Resting membrane potential (RMP) was measured 10 sec after the beginning of whole cell recording (rupture of the membrane under the recording pipette). The dependence of firing rate on the injected current was obtained by injection of current steps (of 500ms duration from 0 to 400 pA in 50 pA increments). Input resistance was calculated from the voltage response to a hyperpolarizing current pulse (−150 pA). SAG ratio was calculated from voltage response -150 pA. The SAG ratio during the hyperpolarizing steps was calculated as [(1-ΔV_SS_/ ΔV _max_) x 100%] as previously reported by (Song, Ehlers, & Moyer, 2015). The membrane time constant was determined using a single exponential fit in the first 100ms of the raising phase of cell response to a 1 second, -150 pA hyperpolarization step.

For measurements of a single action potential (AP), after initial assessment of the current required to induce an AP at 15ms from the start of the current injection with large steps (50 pA), a series of brief depolarizing currents were injected for 10ms in steps of 10 pA increments. The first AP that appeared on the 5ms time point was analyzed. A curve of dV/dt was created for that trace and the 30 V/s point in the rising slope of the AP was considered as threshold (Chakraborty et al., 2017). AP amplitude was measured from the equipotential point of the threshold to the spike peak, whereas AP duration was measured at the point of half-amplitude of the spike. The medium after-hyperpolarization (mAHP) was measured using prolonged (3 seconds), high-amplitude (3 nA) somatic current injections to initiate time-locked AP trains of 50 Hz frequency and duration (10 –50 Hz, 1 or 3 s) in pyramidal cells. These AP trains generated prolonged (20 s) AHPs, the amplitudes and integrals of which increased with the number of APs in the spike train. AHP was measured from the equipotential point of the threshold to the anti-peak of the same spike (Gulledge et al., 2013). Fast (fAHP), and slow AHP (sAHP) measurements were identified as previously described (Andrade et al., 2012; Song and Moyer, 2018). Series resistance, Rin, and membrane capacitance were monitored during the entire experiment. Changes of at least 30% in these parameters were criteria for exclusion of data.

### Classification of Burst and Regular spiking neurons

At the end of recordings, neurons were classified as either burst (BS) or regular spiking (RS) as reported previously (Kim et al., 2015; Song et al., 2015). Briefly, neurons that fired two or more action potentials (doublets or triplets) potential towards a depolarizing current step above the spike threshold current were defined as burst spiking (BS). Regular spiking (RS) neurons on the other hand, were defined as neurons that fired single action potential in response to a depolarizing current step above spike threshold (Extended Figure 1-2A).

### Statistical analysis of individual intrinsic properties across treatments

Individual intrinsic properties of aIC-BLA projecting neurons in the respective treatment groups (Figures 1-4) were analyzed using appropriate statistical tests (One-way or Two-way ANOVA, GraphPad Prism®), as defined in the Statistics Table. Two-way repeated measurements of analysis of variance (RM-ANOVA) followed by Sidak’s (for two groups) or Tukey’s (for more than two groups) post-hoc multiple comparison test was performed for firing properties. The intrinsic properties were determined with Two-tailed unpaired t-tests, and One-way ANOVA followed by Tukey’s or Dunn’s multiple comparisons test were used. For all tests, ∗p < 0.05 was considered significant. Following spike-sorting, the ratio of BS:RS aIC-BLA projecting neurons in the sampled population was compared across our treatments (Mann-Whitney test, GraphPad Prism®). Similarly, individual intrinsic properties in BS and RS aIC-BLA projecting neurons were analyzed following spike-sorting (One-way or Two-way ANOVA, GraphPad Prism®). All data reported as mean ± standard error (SEM).

### Immunohistochemistry

From each electrophysiological recording, three 300μm-thick mouse brain slices were obtained starting from Bregma coordinates 1.78, 1.54 and 1.18, respectively. Slices were washed with PBS and fixed using 4% paraformaldehyde in PBS at 4^0^C for 24 hours. Slices were then transferred to 30% sucrose/PBS solution for 48 hours and mounted on glass slides using Vectashield® mounting medium with DAPI (H-1200). Slides were then visualized using a vertical light microscope at 10x and 20x magnification (Olympus CellSens Dimension®). Images were processed using Image-Pro Plus® V-7 (Media Cybernetics). The localization of labelled mCherry+ neurons in the agranular aIC - where recordings were obtained from, was quantified manually across three Bregma-matched slices, for each animal. Quantification was done using randomly assigned IDs for individual animals, regardless of treatment. Representative images were additionally processed using the Olympus CellSens 2-D deconvolution® function.

### Principal component analysis (PCA) of the profile of intrinsic properties across treatment groups

Principal component analysis (PCA) of the standardized intrinsic properties of the LIV-VI aIC-BLA (Figure 5; Extended Figure 5-1) was performed using the correlation matrix on GraphPad Prism9, MATLAB R2020b, and IBM SPSS Statistics 27. The covariance matrix was used for each PCA was performed in six behavioral groups, the low memory prediction (Saccharin 1x, n=20; Saccharin 2x, n=20, and Extinction, n=14), and the high memory prediction (Saccharin 5x, n=18; CTA retrieval, n=27, and Reinstatement, n=15), RS vs. BS neurons. A total of 114 neurons (BS vs. RS) across all intrinsic properties and excitability changes (50–400 pA) (Extended Figure 5-1A), and later all intrinsic properties with only 350 pA (highest excitability differences between treatment groups; Extended Figure 5-1, B). PCA was conducted on 63 burst spiking neurons using 12 variables: 350 (pA), RMP (mV), mAHP (mV), sAHP (mV), fAHP (mV), IR (MΩ), Sag Ratio, Time constants (msec), AP amplitude (mV), AP Halfwidth (msec), AP threshold (mV), Rheobase (pA), (Figure 5 A &B). The adequacy of the sample was evaluated using the Bartlett’s test and the Kaiser-Meyer-Olkin (KMO) measure was applied. The degrees of freedom (df) were calculated using the following formula:

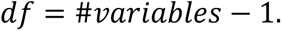

**Figure 5:**
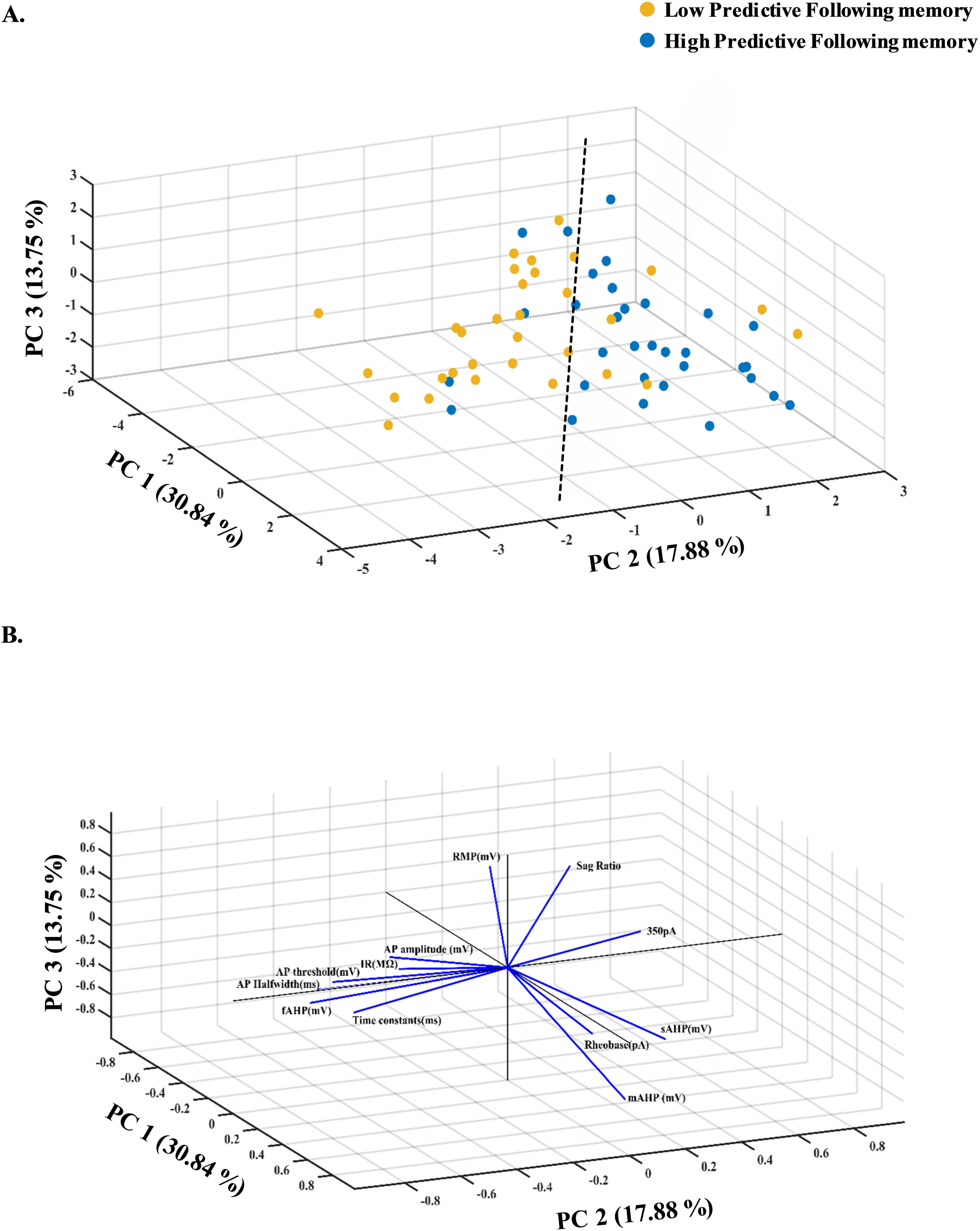
The intrinsic properties of burst-spiking LIV-VI aIC-BLA projecting neurons represent taste experience and the probability for further learning A) Data across all intrinsic properties from BS LIV-VI aIC-BLA neurons of the Saccharin 1x, Saccharin 2x and Extinction groups were combined and assigned to the Low predictive following memory group (32 BS cells). Conversely, the intrinsic properties of BS LIV-VI aIC-BLA neurons from animals having undergone CTA retrieval, 5x Saccharin, and Reinstatement were combined and assigned to the High predictive following memory group (31 BS cells). The resultant three-dimensional scatter representation of the two groups encompassed Excitability at 350pA; AP amplitude, AP halfwidth, AP threshold; fAHP, mAHP, sAHP; IR, Rheobase, RMP, SAG ratio and τ in BS LIV-VI aIC-BLA neurons. B) Three-dimensional representation of the contribution of individual parameters (loadings matrix) to the principal components segregating the two groups of treatments (scores matrix).

The number of principal components was chosen according to the percentage of variance explained (>75%). The parallel analysis evaluated the optimal number of components and selected 3 PCs, explaining 62.47% of the variance. Oblique factor rotation (par) of the first three PCA components, using a standard ‘rotatefactors’ routine from MATLAB Statistics Toolbox. This approach maximizes the varimax criterion using an orthogonal rotation. To optimize variance, oblique factor rotation (paramax) was used, and the threshold chosen to define a variable as a significant contributor was a variance ≥ 0.7 given the small sample size. The correlation matrix was adequate as the null hypothesis of all zero correlation was rejected [χ_66_^2^=387.444, p<0.001], and KMO exceeded 0.5 (KMO=0.580).

To calculate the proportion of the variance of each variable that the principal components can explain, communalities were calculated and ranged from 0.426 to 0.897 (extended Figure 5-1, -A C). The communalities scores were calculated using the following formula: *=* 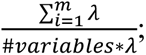 where *m* is the number of selected PCs. The threshold chosen was Comm ≥ 60%.

Due to the imbalance in sample sizes between groups, the PCA space is biased in favor of the group with bigger sample size. The BS neurons in the six behavioral groups previously mentioned were resampled to ensure that sample sizes were balanced across groups datasets (Figure 5, A&B). Particularly, we reduced the number of Saccharin 1X and 2X observations by using random sampling (“randn” function in MATLAB); for Saccharin 1x, we chose 10 of the 17 total elements, and for Saccharin 2x, we selected 10 of the 13 total elements.

### K-means clustering

To assess the distribution of bursting neurons in multidimensional space, we performed a k-means cluster analysis in MATLAB (k = 2 clusters, maximum iterations was 100 with random starting locations, squared Euclidean distance metric used) for the principal components, which explained 62.47% of the variance in intrinsic properties (Figure 5; extended Figure 5-1 and 5-2).

## Results

To prove or refute our hypotheses, we conducted a series of electrophysiological recordings in slice preparation from the mouse aIC, in which we labelled aIC-BLA projecting neurons using retrograde adeno-associated viral tracing – retro AAV (see Methods). Electrophysiological recordings were obtained from the aIC-BLA projecting neurons in LI-III (Table 2) and LIV-VI (Table 1), following novel appetitive or aversive taste stimuli (Figure 1), following appetitive or aversive taste memory retrieval (Figure 2), as well as following extinction and reinstatement (Figures 3-4). We focused on deep-lying LIV-VI aIC-BLA projecting neurons (Table 1), as intrinsic properties in superficial LI-III aIC-BLA projecting neurons were unaffected by taste identity, familiarity, or valence (Table 2). We measured action potential (AP) firing frequency in response to incrementally increasing depolarizing current injections, as well as 11 distinct of intrinsic properties (Tables 1-2 and Statistics Table): resting membrane potential (RMP); slow, medium and fast after-hyperpolarization (sAHP, mAHP, fAHP); input resistance (IR), SAG ratio; the amplitude, half-width and threshold for APs; the time taken for a change in potential to reach 63% of its final value (membrane time constant - τ), as well as the minimum current necessary for AP generation (Rheobase). Statistical analysis was conducted using repeated measures one- or two-way ANOVA (see Statistics table).

### When taste is both appetitive and novel, excitability in LIV-VI aIC-BLA projection neurons is increased

To delineate the mechanisms through which novelty is encoded on the LIV/VI aIC-BLA projection, we labeled the projection (see methods), and compared the intrinsic properties across neutral, innately aversive, and innately appetitive taste stimuli. Following surgery recovery mice were randomly assigned to the following behavioral groups: Water (Control for procedure, Water; n=6 animals, 23 cells), 0.5 % Saccharin for the first (Novel innate appetitive, Saccharin 1x; n=5 animals, 20 cells), or fifth time (Familiar appetitive, Saccharin 5x; n=6 animals, 18 cells), 0.04% Quinine (Novel aversive, Quinine 1x; n=4 animals, 19 cells), or cage controls that did not undergo water-restriction (Base-line control, Cage control; n=4 animals, 19 cells). This approach allowed us to examine excitability changes that relate to the innate aversive/appetitive nature and novelty/familiarity associated with tastants, while accounting for the effects of acute drinking, as well as the water restriction regime itself. Guided by evidence regarding the induction of plasticity cascades, the expression of immediate early genes, as well as the timeframes involved in LTP and LTD at the IC (Rosenblum et al., 1997; Hanamori et al., 1998; Jones et al., 1999; Escobar and Bermúdez-Rattoni, 2000), the five treatment groups were sacrificed 1 hour following taste consumption. Even though changes in activity can be observed within seconds to minutes, depending on their novelty, salience and valence (Barot et al., 2008; Lavi et al., 2018; Wu et al., 2020), sensory experiences can modulate the function of IC neurons for hours (Juárez-Muñoz et al., 2017; Rodríguez-Durán et al., 2017; Haley et al., 2020; Kayyal et al., 2021; Yiannakas et al., 2021).We had previously identified a CaMKII-dependent short-term memory trace at the IC that last for the first 3 hours following taste experiences, regardless of their valence (Adaikkan and Rosenblum, 2015). To address whether similar time-dependency of the physiological correlations engaged by the IC during novel taste learning, a 6^th^ group was sacrificed 4 hours following novel saccharin exposure (Figure 1 – Saccharin 1x (4hrs)).

Daily water intake prior to the final taste exposure and the acquisition of electrophysiological data was not different among the five groups that underwent water restriction (Figure 1B, One-way ANOVA, p=0.4424, F =0.9766, R squared =0.1634). However, excitability in response to incremental depolarizing currents was significantly different between the six groups (Figure 1D – Two-way ANOVA, p<0.0001, F (8,880) =1269). Exposure to saccharin for the first time (i.e., novel appetitive), at the 1-hour time-point, resulted in enhanced excitability on the aIC-BLA projection compared to all other groups (Figure 1D, see Table 1). Conversely, fAHP (Figure 1H; One-way ANOVA, p<0.0001, F =8.380, R squared =0.2758) in the Quinine 1x group was increased compared to all other groups, in contrast to Saccharin 1x where it was most suppressed (see Table 1). In fact, fAHP in the Saccharin groups recorded at 1hr (p<0.0001, z =5.150) or 4hours (p=0.0099, z =3.406) following novel taste consumption was suppressed compared to innately aversive Quinine 1x (Figure 1H). Even though fAHP in the Saccharin 1x group was suppressed compared to both the Cage control (p=0.0136, z =3.318) and Water (p=0.0177, z =3.243) groups, this was not the case for the Saccharin 1x (4hr) group (p>0.9999 for both – see Table 1). Importantly, fAHP (Figure 1H) was nearly identical in treatment groups where the tastant could be deemed as highly familiar and safe, such as the Cage control group (that did not undergo water restriction), as well as animals in the Water or Saccharin 5x groups (that had undergone water restriction).

Significant differences in terms of τ (Figure 1J; Kruskal-Wallis test, p=0.0005, Kruskal-Wallis statistic=21.91), were only observed between the Cage control and Saccharin 1x (4hrs) groups (p=0.0040, z=3.647), as well as the Saccharin 1x and Saccharin 1x (4hrs) groups (p=0.0029, z=3.725). On the other hand, significant differences in AP half-width (Figure 1I; Kruskal-Wallis test; p=0.0125, Kruskal-Wallis statistic=14.54) were only observed between the Saccharin 1x (4hrs) compared to Saccharin 1x (p=0.0065, z =3.519) groups (see Table 1).

These results demonstrate that in the context of taste novelty, innately appetitive saccharin drove increases in excitability and decreases in fAHP of LIV-VI aIC-BLA projecting neurons, compared to innately aversive quinine (Figure 1D, H). Compared to the Cage control and Water groups, fAHP on the projection was significantly enhanced by innately aversive quinine and was suppressed by innately appetitive novel saccharin (Figure 1H). However, the effect of appetitive taste novelty on firing frequency was time-dependent, as it was observed at 1hr, but not 4hrs following novel taste exposure (Figure 1D). Furthermore, following familiarity acquisition for saccharin (Saccharin 5x), excitability was suppressed compared to Saccharin 1x, matching the Cage control, Water, and Quinine 1x groups (Figure 1D). This led us to consider whether increased excitability is not related to taste identity or palatability (Wang et al., 2018), but the perceived salience of taste experiences, which encompasses both novelty and valence (Ventura et al., 2007; Kargl et al., 2020). Previous studies have suggested that the induction of plasticity signaling cascades and IEGs in pyramidal neurons of the aIC (commonly used as surrogates for changes in excitability), is a crucial step for the association of taste and visceral information during CTA learning (Adaikkan and Rosenblum, 2015; Soto et al., 2017; Wu et al., 2020). Activation of the aIC-BLA projection is indeed necessary for the expression of neophobia towards saccharin (Kayyal et al., 2021), as well as for CTA learning and retrieval (Kayyal et al., 2019). Yet, its chemogenetic inhibition does not affect the attenuation of neophobia, nor the expression of aversion towards innately aversive quinine (Kayyal et al., 2019). Furthermore, aversive taste memory retrieval necessitates increases in pre-synaptic inhibitory input on the projection (Yiannakas et al., 2021). Bearing this in mind, we hypothesized that increases in excitability on the projection could be indicative of a labile state of the taste trace at the aIC, which manifests when taste cues are not (yet) highly predictive of the visceral outcome of the sensory experience (Bekisz et al., 2010; Galliano et al., 2021). In such a scenario, taste memory retrieval following strong single-trial aversive learning would be expected to result in decreased excitability compared to control animals. To assess this hypothesis, we next examined intrinsic excitability in mice retrieving an appetitive (Saccharin 2x, CTA retrieval control) or learned aversive memory (CTA retrieval) for saccharin.

### Learned aversive taste memory retrieval suppresses the excitability of LIV-VI aIC-BLA projecting neurons

Following recovery from rAAV injection, mice in the CTA retrieval group underwent water restriction and CTA conditioning for 0.5% saccharin (see Methods, Figure 2A). Electrophysiological recordings were obtained from aIC-BLA neurons 3 days later, 1 hour following retrieval (n=8 animals, 27 cells). Mice in the Saccharin 2x group on the other hand, were familiarized with saccharin without conditioning, and recordings were obtained within the same period, following retrieval (n=5 animals, 20 cells). Through this approach we aimed to examine the hypothesis that like innately aversive and highly familiar appetitive responses (Figure 1), learned aversive taste memory retrieval would be correlated with suppression of the intrinsic excitability on the projection.

As expected, CTA retrieval mice, exhibited suppressed consumption of the conditioned tastant compared to control animals that were only familiarized with saccharin (Figure 2B, Mann-Whitney test, p=0.0085; Sum of ranks: 52.50, 38.50; Mann-Whitney U =2.500). Intrinsic excitability in LIV-VI aIC-BLA projecting neurons was increased in response to depolarizing current injections (Figure 2D; p<0.0001, F (8, 360) =483.3), and was significantly different between the two treatments (p=0.0014, F (1, 45) =11.60). Excitability was enhanced in the Saccharin 2x group compared to CTA retrieval, while a significant interaction was identified between the treatment and current injection factors (p<0.0001, F (8, 360) =9.398). Fast AHP (fAHP) on LIV-VI aIC-BLA projecting neurons tended to be increased in the CTA retrieval group (see Table 1), however differences compared to Saccharin 2x failed to reach significance (Unpaired t-test; p=0.0527, t=1.990, df=45). Conversely, AP amplitude in the Saccharin 2x group was significantly decreased compared to CTA retrieval (Figure 2G; Unpaired t-test; p=0.0002, t=3.983, df=45). In addition, the CTA retrieval group exhibited significantly decreased IR (Figure 2H; Unpaired t-test; p=0.0036, t=3.072, df=45) and significantly enhanced SAG ratio (Figure 2I; Unpaired t-test; p=0.0037, t=3.060, df=45), compared to Saccharin 2x. In accord with our hypothesis, excitability on LIV-VI aIC-BLA projecting neurons was suppressed by aversive taste memory retrieval. We have previously shown that compared to CTA retrieval and reinstatement, appetitive memory retrieval and extinction were associated with (a) an enhancement of IEG induction (c-fos and Npas4) at the aIC, and (b) decreased frequency of pre-synaptic inhibition on the aIC-BLA (Yiannakas et al., 2021). In accord, other published work investigating the induction of IEG in the rodent IC, found that consistent with a reduction in spiking activity (Grossman et al., 2008), the induction of c-fos at the IC was suppressed by aversive taste memory retrieval (Haley et al., 2020). Earlier studies have also reported increases in c-fos following the extinction of cyclosporine A-induced CTA (Hadamitzky et al., 2015). We thus hypothesized that if excitability in these cells serves as key node for a change in valence prediction, extinction - which constitutes a form of appetitive re-learning, would be associated with enhanced excitability compared to CTA retrieval and reinstatement (Berman, 2003; Suzuki et al., 2004; Morrison et al., 2016; Slouzkey and Maroun, 2016). In addition, through these extinction and reinstatement studies, we were able to examine the real-life relevance of these changes on intrinsic excitability, in a context where behavioral performance reflects the balance between contrasting memories and the availability of retrieval cues (Figure 3).

### The predictability of the valence arising from taste experiences determines the profile of intrinsic properties of LIV-VI aIC-BLA projecting neurons

Using similar approaches, electrophysiological recordings were obtained from LIV-VI aIC-BLA projecting neurons from mice having undergone unreinforced CTA extinction (Extinction; n=5, 14 cells), or US-mediated CTA reinstatement (Reinstatement; n=3 animals, 15 cells). Behaviorally, the two groups of animals were similar in terms of their aversion profile over 9 unreinforced extinction sessions (Figure 3B; 2-way ANOVA; Extinction: p<0.0001, F (8, 54) =13.44; Treatment: p=0.0681, F (1, 54) =3.466; Interaction: p=0.9697, F (8, 54) =0.2803). As expected, saccharin consumption during the test day in the Reinstatement group was decreased compared to Extinction (Figure 3C; Mann-Whitney test; p=0.0179; Sum or ranks: 30, 6; Mann-Whitney U=0). Consistent with our findings in Figure 2, aversive taste memory retrieval in the Reinstatement group was associated with suppressed excitability compared to the Extinction group (Figure 3E; 2-way ANOVA, Current injection: p<0.0001, F (8, 216) =370.1; Treatment: p=0.0297, F (1, 27) =5.291; Interaction: p<0.0001, F (8, 216) =10.30). CTA Reinstatement was also associated with increases in the AP threshold (Figure 3F; Unpaired t-test: p=0.0076, t=2.887, df=27) and τ (Figure 3G; Unpaired t-test: p=0.0153, t=2.589, df=27) compared to Extinction.

Unlike animals that underwent familiarization with the tastant without conditioning (Figure 1), excitability on the projection in the Extinction group was not suppressed by familiarization (Figure 3). Conversely, even though the intrinsic mechanisms employed would appear to differ, aversive taste memory retrieval regardless of prior experience, was associated with baseline excitability of the aIC-BLA projection (Figure 3). Our findings in this section (Figure 3), revealed that during taste memory retrieval, excitability on the projection is not solely dependent on the relevant novelty or appetitive nature of tastants, and does not subserve the persistence of CTA memories (Figure 2). Instead, excitability on the aIC-BLA projection is indeed shaped by prior experience but is best predicted by the probability for further aversive (re)learning.

Next, to distinguish between intrinsic properties changes that reproducibly reflect taste identity, familiarity, and valence over the course of time and experience, we compared the profile of intrinsic properties across pairs of behavioral groups in which the currently perceived novelty, as well as innate or learned valence associated with taste was notably different. Through this comparison, we were led to conclude that excitability on aIC-BLA projecting neurons is driven by taste stimuli of positive valence, however this effect is dependent on subjective experience and the possibility for further associative learning (Figure 4A). Excitability on aIC-BLA projecting neurons in the treatment groups where the tastant was perceived as appetitive (Saccharin 1x, Saccharin 2x and Extinction), was closely matched, and was significantly enhanced compared to the innately or learned aversive (Quinine 1x, CTA retrieval and Reinstatement) groups (Figure 4A; Two-way ANOVA; Current injection: p<0.0001, F (5, 872) =1218; Treatment: p=0.0014, F (5, 109) =4.281; Interaction: p<0.0001, F(40, 872) =4.978). As previously identified in Figure 1H, fAHP reflected the innate aversive nature of the tastant, being increased in the Quinine 1x group compared to all other groups (Figure 4B; One-way ANOVA; F =10.65, p<0.0001, R squared =0.3283, see Table 1). Significant differences in IR (Figure 4C; One-way ANOVA; F = 2.775, p=0.0213, R squared =0.1129) and SAG ratio (Figure 4D; One-way ANOVA; F =2.610, p=0.0286, R squared =0.1069) were only observed between the CTA retrieval and Saccharin 2x groups. AP amplitude (Figure 4E; One-way ANOVA, p=0.0054, F=3.526, R squared =0.1392) in the Saccharin 2x group was suppressed compared to both CTA retrieval (p=0.0129, q=4.768, df=109) and Quinine 1x (p=0.0087, q=4.944, df=109). Conversely, the Extinction and Reinstatement groups, where familiarity with the tastant was the highest, exhibited increased AP half-width compared to all other groups (Figure 4F; One-way ANOVA, Kruskal-Wallis test; p=0.0002; Kruskal-Wallis statistic, 24.03). Significant differences in terms of τ (Figure 4G; One-way ANOVA, p=0.0047, F =3.606) were only observed in comparing the Saccharin 1x and Reinstatement groups (p=0.0022, q=5.525, df=109). Hence, neuronal excitability is indeed a feature associated with predictive power to modulate taste valence, however it does not fully reflect the breadth of intrinsic property changes among the different behavioral groups.

### The predictability of taste valence intrinsic is primarily reflected on the excitability of burst-spiking, but not regular-spiking LIV-VI aIC-BLA projecting neurons

Our initial analysis of individual intrinsic properties (Figures 1-3) highlighted that excitability is enhanced following appetitive experiences in which the internal representation is still labile and is associated with the possibility for further aversive learning (novelty or extinction). Conversely, following extensive familiarization, aversive conditioning, or reinstatement, whereby taste exposure leads to memory retrieval of specific valence, excitability on LIV-VI aIC-BLA projecting neurons was similar to baseline (Figures 1 and 4). While the precise mechanism through which sensory input is encoded at the cortex (and other key regions), is still a matter of ongoing research, studies indicate that bursting in cortical layer V pyramidal neurons can encode oscillating currents into a pattern that can be reliably transmitted to distant post-synaptic terminals (Kepecs and Lisman, 2003; Samengo et al., 2013; Zeldenrust et al., 2018). Spike burst is defined as the occurrence of three or more spikes from a single neurons with <8ms intervals (Ranck, 1973; Connors et al., 1982). In brain slices from naïve mice, half of the neurons of a given structure exhibit burst firing, while the distribution of burst-spiking (BS) to regular-spiking (RS) neurons, changes along the anterior-posterior axis of the subiculum (Staff et al., 2000; Jarsky et al., 2008). Importantly, the two cell types fine-tune the output of brain structures by virtue of differences in synaptic plasticity, as well as intrinsic excitability mechanisms (Graves et al., 2012; Song et al., 2012). Furthermore, there are changes in the ratio of BS:RS neurons in individual brain structures, as well as differences in the recruitment of signaling events, ion channels and metabotropic receptors among the two cell types (Wozny et al., 2008; Shor et al., 2009). Correspondingly, complex region and task-specific rules govern the molecular and electrophysiological mechanisms through which information encoding and retrieval takes place in the two cell types (Dunn et al., 2018; Dunn and Kaczorowski, 2019). Little is currently known regarding the influence of cell identity in the repertoire of plasticity mechanisms employed by the IC to facilitate taste-guided behaviors (Maffei et al., 2012; Haley and Maffei, 2018).

Our post-hoc spike sorting analysis allowed us to distinguish between BS and RS LIV-VI aIC-BLA projecting neurons, and thus their relative contribution to behaviorally driven changes in the suit of intrinsic properties (Extended Figures 1-3, 2-1 and 3-1). Through this comparison, we uncovered that Saccharin 1x differed to other groups in terms of excitability and fAHP in BS LIV-VI aIC-BLA neurons (Extended Figure 1-3), while no such changes were observed in RS neurons (see Summary of RS intrinsic properties table no.4). Similarly, excitability in the Saccharin 2x group was significantly enhanced compared to CTA retrieval in BS-, but not in RS LIV-VI aIC-BLA neurons (Extended Figure 2-1A, F). Significant differences in IR, SAG ratio and AP amplitude between CTA retrieval and Saccharin 2x were primarily driven by BS LIV-VI aIC-BLA neurons (Extended Figure 2-1B-D, G-I). Conversely, significant differences in AP half-width between the aversive and appetitive memory retrieval groups were only observed in RS neurons (Extended Figure 2-1J). Correspondingly, excitability in the Extinction group was enhanced compared to Reinstatement in BS-, and not RS LIV-VI aIC-BLA neurons (Extended Figure 3-1A, H). Indeed, excitability on BS LIV-VI aIC-BLA neurons following extinction and reinstatement, reflected the subjective predictability of taste memory retrieval, being high following extinction compared to reinstatement (Extended Figure 3-1). However, this effect was mediated through alternative mechanisms compared to single-trial learning and memory retrieval (Extended Figure 1-3, 2-1, 3-1). Significant differences between the Extinction and Reinstatement groups, were observed in terms of the sAHP, AP threshold, SAG ratio and τ in BS but not in RS LIV-VI aIC-BLA neurons (Extended Figure 3-1B-F).

Encouraged by these findings, we focused on the Saccharin 1x, Saccharin 2x, Saccharin 5x, CTA Retrieval, Extinction and Reinstatement groups, as to isolate the contribution of BS LIV-VI aIC-BLA neurons in encoding the subjective predictability of taste experience during taste learning, re-learning, and memory retrieval (Figure 5). Consistent with studies in the hippocampus (Graves et al., 2016), we found that the percentage of BS LIV-VI aIC-BLA projecting neurons in the sampled population was highest in the context of novel taste learning (Extended Figure 1-2B: Saccharin 1x, 85%), and subsided following progressive familiarization (Extended Figure 1-2B; Saccharin 2x, 65%, Mann-Whitney test: p=0.0562; Sum of ranks: 303.5, 161.5; Mann-Whitney U =70.50; Saccharin 5x, 55.56%, Mann-Whitney test: p=0.0034; Sum of ranks: 291, 87; Mann-Whitney U =32;). Interestingly, animals retrieving CTA, exhibited the lowest proportion of BS neurons among the six treatments (Extended Figure 1-2B; CTA retrieval, 44.44% BS), and significant differences were observed compared to control animals (Extended Figure 1-2B; Saccharin 2x, 65% BS; Mann-Whitney test: p=0.0102; Sum of ranks: 257, 208; Mann-Whitney U =55). Thus, the ratio of BS:RS LIV-VI aIC-BLA projecting neurons was plastic in relation to experience and was highest in response to appetitive novelty – in accord with studies investigating the intrinsic excitability of subiculum output neurons in relation to contextual novelty and valence encoding (Dunn et al., 2018). Indeed, the ratio of BS:RS LIV-VI aIC-BLA neurons was progressively suppressed by familiarity acquisition (Saccharin 1x> 2x> 5x), as well as following aversive taste memory recall (CTA retrieval), compared to both appetitive learning (Extended Figure 1-2B; Saccharin 1x, Mann-Whitney test: p<0.0001; Sum of ranks: 407, 188; Mann-Whitney U =35) and re-learning (Extended Figure 1-2B; Extinction, Mann-Whitney test: p=0.0007; Sum of ranks: 182, 224; Mann-Whitney U =29). However, our comparison failed to account for the influence of complex experiences over time, as differences between Extinction and Reinstatement failed to reach significance (Extended Figure 1-2B; Mann-Whitney test: p=0.3870, Sum of ranks: 133.5, 97.50, Mann-Whitney U =42.50), while perplexingly, the ratio of BS:RS aIC-BLA neurons in these groups was differentially increased compared to Saccharin 5x (Extended Figure 1-2B; Extinction, Mann-Whitney test: p=0.0300; Sum of ranks: 81, 150; Mann-Whitney U =26; Reinstatement, Mann-Whitney test: p=0.3498; Sum of ranks: 90, 120; Mann-Whitney U =35;), but not Saccharin 2x (Extended Figure 1-2B; Extinction, Mann-Whitney test: p=0.2397; Sum of ranks: 143.5, 156.5; Mann-Whitney U =52.50; Reinstatement, Mann-Whitney test: p>0.9999; Sum of ranks: 153.50, 122.5; Mann-Whitney U =62.50). No further statistics were performed in intrinsic properties of aIC-BLA regular spiking neurons representing (Figure 1 and Figure 3), because of the small sample size.

Changes in the intrinsic properties of neuronal ensembles have recently been suggested to contribute to homeostatic mechanisms integrating both cellular and synaptic information (Wu et al., 2021). In our current study, we randomly sampled from neuroanatomically defined LIV-VI aIC-BLA projecting neurons, and even following spike sorting (Extended Figures 1-2), the probability of recording from engram cells (10% of neurons within a region) would be extremely low (Tonegawa et al., 2015). Importantly, the correlative nature does not exclude the possibility that these changes are the consequence of representational drift (Driscoll et al., 2017). We thus set out to examine the hypothesis that applying linear dimension reduction method on the complement of intrinsic properties recorded in BS LIV-VI aIC-BLA neurons would allow us to distinguish between taste experiences that differ in terms of their perceived predictability (or the associated probability for further aversive learning).

### Principal component analysis of the profile of intrinsic properties in BS LIV-VI aIC-BLA projecting neurons separates treatment groups in relation to the perceived predictability of taste valence for saccharin

We assigned six of our treatment groups into highly predictive scenarios (Saccharin 5x, CTA retrieval and Reinstatement), and low predictive scenarios (Saccharin 1x, Saccharin 2x, Extinction). We used parallel analysis to select the components across the complement of intrinsic properties in each treatment group, with the first three principal components (PC1-3) explaining 30.84%, 17.88%, and 13.75% of the total variance, respectively, and 62.47% of the variance, collectively (Figure 5-2). PC1 (Figure 5B, Extended Figure 5-2) was characterized by strong negative loadings for Rheobase (−0.88304), sAHP (−0.85985) and mAHP (−0.82421), while a positive correlation was identified for IR (0.694764). The direction of PC2 (Figure 5B, Extended Figure 5-2) was positively correlated with fAHP (0.682681) and AP halfwidth (0.614103) and was negatively correlated with Excitability at 350pA (−0.69587). PC3 (Figure 5B, Extended Figure 5-2) positively correlated with SAG ratio (0.889949) and RMP (0.682681), whereas a significant negative correlation with mAHP (−0.6095) was also identified. Unlike aversive or appetitive taste memory retrieval (i.e., highly predictive), appetitive novelty or extinction learning (i.e., low predictive), was associated with increased input resistance, faster action potential generation and suppressed afterhyperpolarization on BS aIC-BLA neurons (Figure 5). Importantly, PCA of the intrinsic properties of LIV-VI aIC-BLA projecting neurons regardless of cell type (BS and RS together, Extended Figure 5-1), failed to segregate the two groups of treatments. This cell-type specific profile of intrinsic properties could provide the framework through which BS LIV-VI aIC-BLA projecting neurons are able to inspect the gastrointestinal consequences associated with tastants over prolonged timescales, when these consequences are not accurately predicted by sensory experience or memory retrieval alone (Adaikkan and Rosenblum, 2015; Lavi et al., 2018; Kayyal et al., 2019).

## Discussion

Learning and memory are subserved by plasticity in both synapse strength and neuronal intrinsic properties (Citri and Malenka, 2008; Sehgal et al., 2013). While Hebbian rules can explain associative learning paradigms where seconds separate the CS and US (Krabbe et al., 2018), additional cellular-level mechanisms are needed to explain how learning operates in paradigms where the time between CS and US extends to hours (Adaikkan and Rosenblum, 2015; Wu et al., 2021). In this study, we demonstrate that following taste experiences, taste percept and prior experience are integrated in the intrinsic properties of the aIC in a time-dependent and cell-type specific manner. We further show that regardless of the identity or prior history associated with taste, the intrinsic properties of BS LIV-VI aIC-BLA projecting neurons encodes the perceived confidence of taste valence attribution.

We focused on the aIC-BLA projection; a circuit causally involved in the acquisition and retrieval of CTA memories (Kayyal et al., 2019, 2021). (Kayyal et al., 2019, 2021). We examined the hypothesis that excitability in aIC-BLA neurons can serve as a taste valence updating mechanism enabling the prolonged ISI between CS and US in CTA learning (Adaikkan and Rosenblum, 2015), and/or contributes to anticipatory valence attribution (Barrett and Simmons, 2015). Our basic proposition diverged from Hebb’s famous postulate that cells with increased excitability over hours can potentially wire together with cells conveying incoming modified valence information (Hebb, 1961).

The confidence with which taste valence is encoded is the product of the subjectively perceived- (a) appetitive or aversive nature of tastants and (b) novelty or familiarity associated with tastants (Russell, 1980; Kahnt and Tobler, 2017). We first examined each axis separately and later in tandem as to better simulate real-life scenarios. We measured the intrinsic properties of aIC-BLA neurons 1 hour following taste experience – a previously established suitable time point (Jones et al., 1999; Haley et al., 2020).

To dissociate novelty-related changes from those involving hydration, taste identity and familiarity; we compared the intrinsic properties of aIC-BLA neurons following Water – a neutral and familiar tastant, Saccharin – an innately appetitive tastant, in the context of novelty (1x) or familiarity (5x), and Quinine – an innately aversive novel tastant (Figure 1). Excitability was high following novel saccharin exposure, but not in response to Quinine (Figure 1D). Indeed, concerted activity at the aIC and BLA encodes the presence, identity and palatability of taste experiences within the 2 seconds preceding swallowing (Katz et al., 2001; Grossman et al., 2008; Fontanini et al., 2009). Palatability can be enhanced as a function of experience (Austen et al., 2016), but can also be suppressed by sensory satiety and alliesthenia (Rolls et al., 1981; Yeomans, 1998; Siemian et al., 2021). However, excitability on the projection was enhanced in response to novelty and was suppressed following familiarization (Figure 1). Further inconsistent with palatability encoding; changes in excitability captured 1 hour following novel saccharin exposure subsided 4 hours later (Figure 1), while excitability remained plastic even following longer periods of water restriction, that could be considered monotonous (Figure 5). Deciphering whether and how aIC-BLA neurons contribute to palatability processing would require *in vivo* recordings to capture taste-evoked changes, within timescales that are beyond the scope and means of our current study (Vincis and Fontanini, 2016).

The correlation identified between excitability and innate current taste valence, encouraged us to examine the predictability of future outcomes following aversive taste memory retrieval. Bearing in mind our previous findings using transcription-dependent activity markers at the aIC (Yiannakas et al., 2021), we hypothesized that aversive taste memory retrieval (CTA retrieval or Reinstatement), would be associated with suppressed excitability compared to stimulus- and familiarity-matched controls (Saccharin 2x and Extinction). Indeed, excitability on aIC-BLA projecting neurons following CTA retrieval was suppressed compared to Saccharin 2x (Figure 2B), while Reinstatement, was also associated with decreased excitability compared to Extinction (Figure 3E). Hence, regardless of the complexity of taste memory retrieval, excitability in aIC-BLA neurons was best expected by the subjective predictability of taste valence - increasing in response to innately appetitive taste experiences in which the perceived possibility for avoidance learning was high/taste valence predictability was low (Figure 4). Conversely, when the subjective confidence with which taste valence was predicted was high, excitability on the projection remained unchanged (Figures 1, 4).

Innately and learned aversive tastants were both associated with suppressed excitability on aIC-BLA projecting neurons compared to appetitive controls, however these effects were mediated through alternative mechanisms (Figures 2-4). Quinine increased fAHP on the projection compared to saccharin, regardless of familiarity or perceived valence (Figures 1 and 4). Post-spike after-hyperpolarization (AHP) has a key function in transducing the summed result of processed synaptic input, directly impacting neuronal excitability in relation to both learning and aging (Oh and Disterhoft, 2020). In pyramidal cells of the hippocampus and cortex, differences in fAHP are mediated by the Ca^2+-^ and voltage-dependent BK currents that promote repolarization at the beginning APs trains (Shao et al., 1999). Interestingly, studies in the prefrontal cortex (PFC), have shown that fear conditioning decreases excitability, whereas extinction training enhances excitability and decreases medium- and slow AHP (Santini et al., 2008; Maglio et al., 2021). At the IC, chronic ethanol consumption has shown to decrease excitability and to increase AHP (Luo et al., 2021). Conversely, oxytocin-dependent signaling at the IC has been shown to promote social affective behaviors, via increases in excitability and decreases in sAHP (Rogers-Carter et al., 2018). Further studies would be necessary to fully address this, but our findings could indicate that enhanced fAHP is induced by innately aversive tastants or quinine specifically.

Unlike Quinine, CTA memory retrieval, was associated with increased AP amplitude and SAG ratio, as well as decreased IR in BS LIV-VI aIC-BLA projecting neurons, compared to control animals (Figure 2, Extended Figure 2-1). On the other hand, the suppressed excitability in the Reinstatement group compared to Extinction was characterized by decreased AP threshold, increased τ, and decreased sAHP (Figures 3, 4, Extended Figure 3-1). The hyperpolarization-activated, cyclic nucleotide-gated current (I_h_) regulates membrane depolarization following hyperpolarization (Hogan and Poroli, 2008; Shah, 2014). The opening of HCN channels generates an inward current, that modulates AHP, RMP and IR in cortical pyramidal and PV interneurons (Yang et al., 2018). However, conductance through I_h_ channels, regulates synaptic integration at the soma of pyramidal neurons, by suppressing excitability, decreasing IR, and increasing τ (Hogan and Poroli, 2008). Evidence indicates that this dichotomous impact of HCN channels on neuronal excitability, is mediated by A-type K channels at the dendrites (Mishra and Narayanan, 2015), and M-type channels at the soma (Hu et al., 2007). Notably, AP half-width was significantly increased in the Extinction and Reinstatement groups that had undergone extinction training, compared to all other saccharin-treated groups (Figure 4F). Mechanistically, this effect could reflect changes in the distribution and/or the properties of voltage- or calcium-gated ion channels (Faber and Sah, 2002; Grubb and Burrone, 2010; Kuba et al., 2010). Such broadening of spike width has also been reported in infralimbic PFC neurons projecting to the amygdala in response to extinction training (Senn et al., 2014). PV-dependent restriction of excitability and burst firing, is instrumental in experience-dependent plasticity in the amygdala (Morrison et al., 2016), the hippocampus (Donato et al., 2013; Xia et al., 2017) and visual cortex (Yazaki-Sugiyama et al., 2009; Kuhlman et al., 2013). Conversely, in the striatum, PV interneurons restrict bursting, calcium influx, and synaptic plasticity to appropriate temporal windows that facilitate learning, but not retrieval (Owen et al., 2018). Elegant recent studies report that rapid eye movement sleep is associated with a PV-dependent somatodendritic decoupling in pyramidal neurons of the PFC (Aime et al., 2022). At the IC, the maturation of GABAergic PV circuits is key for multisensory integration and pruning of cross-modal input to coordinate the detection of relevant information (Gogolla et al., 2014). Activation of IC PV disrupts the expression of taste-guided goal-directed behavior (Vincis et al., 2020), and enhances taste-guided aversive responses (Yiannakas et al., 2021). Our findings could be indicative of a prediction-dependent decoupling mechanism at the IC, whereby the restriction of bursting activity in LIV-VI aIC-BLA neurons impinges on innate drives towards the tastant and further learning, depending on prior experience.

We further probed our results and hypotheses using PCA and attempted to segregate behaviors based on the perceived ability of the CS to predict the consequences of sensory experience, and the probability for further learning (Figure 5). We focused on BS LIV-VI aIC-BLA projecting neurons since bursting has been implicated in coincidence detection by deep-layer neurons (Boudewijns et al., 2013; Shai et al., 2015), as well as the encoding of novelty and valence relating to different modalities (Song et al., 2015; Dunn et al., 2018; Yousuf et al., 2019). Our PCA of intrinsic properties in BS LVI-VI aIC-BLA projecting neurons demonstrated that distinct plasticity rules are at play depending on the balance between the probability for associative learning and the certainty with which taste predicts the valence of experience during retrieval (Figure 6). We propose that increased excitability and reduced fAHP on BS LIV-VI aIC-BLA projecting neurons might represent a transient neuronal state in the absence of adequate predictive cues for the outcome of taste experiences (Adaikkan and Rosenblum, 2015).

**Figure 6:**
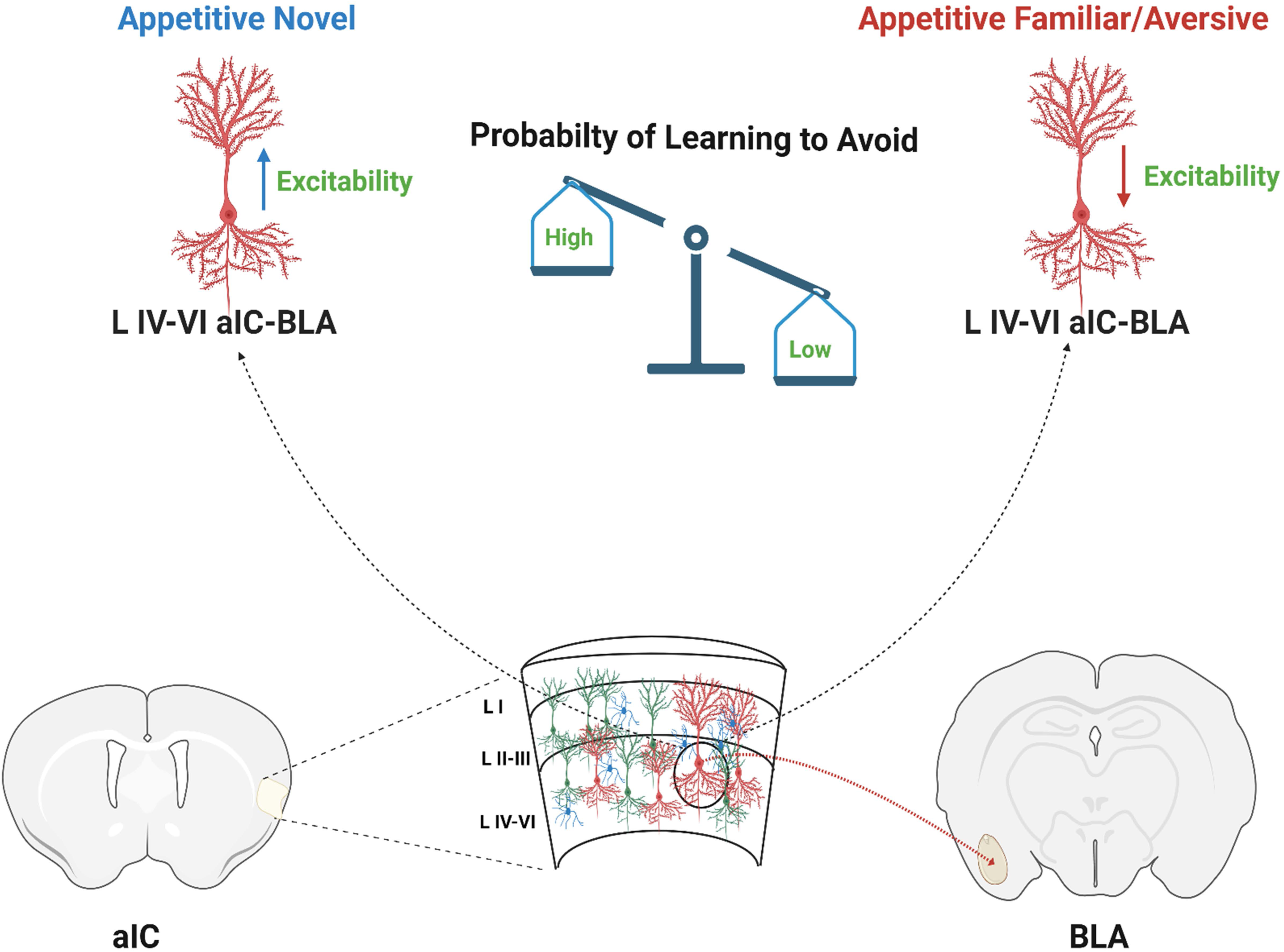
A model for excitability changes in IV-VI aIC-BLA neurons when the taste salience is low and highly predictive following the taste memory

- The unpredictability of future valence outcomes: Novel appetitive taste experiences /extinction of previously learned aversive taste experiences, increases the excitability (blue arrow) of aIC-BLA projecting neurons (red). This increased excitability is specific to the deep-layer aIC-BLA projecting neurons (LIV-VI).
- When the taste valence is highly predictable (familiar/aversive), excitability is reduced in LIV-VI aIC-BLA projecting neurons (red arrow).
- LIV-VI aIC-BLA neurons remain excitable facilitating the association of the taste memory trace with visceral pain when the stimulus is not adequately predictive of the outcome of the sensory experience.

The IC has long been considered crucial for interoception, which is increasingly understood to be supported by distinct direct or indirect functional bidirectional connectivity. Indeed, interoceptive inputs relating to the processing, or anticipation of physiological states of hydration and satiety manifest at the pIC (Livneh et al., 2017, 2020; Livneh and Andermann, 2021). However, this is rarely the case when it comes to physiological hydration or satiety inputs and the aIC, that has is primarily involved in interoceptive processes in the context food poisoning or CTA (Chen et al., 2020; Wu et al., 2020). As other studies currently in press demonstrate, hydration correlates and requires suppressed activity in aIC-BLA and increases in pIC-BLA CB1 receptor-expressing neurons (Zhao et al., 2020; Nicolas et al., 2022). Under uncertain conditions that are associated with greater potential significance, recruitment of the aIC is thought to contribute to attention, effort, and accurate processing (Lovero et al., 2009), as to identify better response options (Preuschoff et al., 2008). In agreement with earlier computational models of the cortical connectivity (Mumford, 1991, 1992), recent work indicates that the aIC facilitates prediction-related encoding driven by hedonics, rather than homeostatic needs (Darevsky et al., 2019; Price et al., 2019; Darevsky and Hopf, 2020). Our results, propose a cellular framework for such an emotional predictive function at the aIC. Future studies will further explore whether and how the interplay between such distinct mechanisms at the aIC, enables its complex role in learning, memory, and decision-making.

## Data availability

All data generated or analyzed during this study are included in the manuscript and supporting files. Source data files have been provided for all figures.

## Author contributions

SKC and AY led the project. AY, SKC and KR designed the research. KR supervised the research. SKC, AY, HK, and MK performed the research. SKC, AY, LM, RS, FC, and SS analyzed the data. AY, SKC and KR drafted the paper. All authors reviewed and contributed to the manuscript.

## Acknowledgments

The authors would like to thank all current members of the Rosenblum labs for their help and support, to the veterinary team headed by Barak Carmi and Corina Dollingher and technical team headed by Yair Bellehsen. Graphical illustrations were created using BioRender.com.

## Extended figures and Tables

Extended Figure 1-1: Histological verification of rAAV-mCherry virus expression and locations of whole-cell patch clamp recordings

A) A representative image showing the distribution of retrograde injections into the BLA and aIC-BLA projection neuron at aIC.

B) Mean localization of BLA projecting neurons of the agranular aIC used for electrophysiological whole cell recordings.

Extended Figure 1-2: The ratio of burst-spiking and regular spiking LIV-VI aIC-BLA projecting neurons changes in relation to the uncertainty associated with taste experiences

A) Representative traces from Burst (BS) and Regular (RS) spiking LIV-VI aIC-BLA projecting neurons in response to rheobase current injections. The neurons showing doublets or triplets in response to rheobase current injection were considered BS. The neurons showing single spike in response to rheobase current injection considered RS. Scale bars 20 mV and 100 msec.

B) Pie charts showing the change in the ratio of BS vs RS LIV-VI aIC-BLA projection neurons, expressed as a percentage of the sampled population across the Saccharin 1x, Saccharin 2x, Saccharin 5x, CTA retrieval, Extinction, and Reinstatement groups.

C) Heat map summary of the change in the ratio of BS vs RS LIV-VI aIC-BLA projection neurons, expressed as a percentage of the sampled population across the six treatment groups.

Extended Figure 1-3: Appetitive novel taste alters the intrinsic properties of burst spiking LIV-VI aIC-BLA neurons

We compared the intrinsic properties of BS and RS LIV-VI aIC-BLA neurons among the Cage control (n=13 cells), Water (n= 11cells), Saccharin 1X (n=17 cells), Quinine 1x (n=9 cells), Saccharin 5x (n=10 cells) and Saccharin 1x (4hrs, n=6 cells).

A) Excitability in BS LIV-VI aIC-BLA was not significantly different among the treatment groups. Two-Way repeated measures ANOVA, Current x Treatment: p<0.0001, Group interaction p= 0.0666.

B) fAHP was significantly enhanced in Quinine 1x (13.67 ±2.681 mV) and Saccharin 5x (11.30 ±1.727 mV) BS neurons compared to Saccharin 1x BS neurons (2.870 ±1.044 mV). One-Way ANOVA, P= 0.0004.

C) Action potential amplitude was significantly different between the groups. Cage controls (56.27 ±1.147 mV), Water (54.21 ±1.572 mV), Saccharin 1x (51.64 ± 1.473 mV), Quinine 1X (58.86 ± 2.003 mV), Saccharin 5x (58.40 ± 1.812 mV), and Saccharin 1x (4hr) (46.79 ± 4.359 mV). One-Way ANOVA, P= 0.0097.

D) Action potential half-width in BS LIV-VI aIC-BLA neurons of the Saccharin 1x (4 hrs.) group (0.8850 ± 0.05943ms) was increased compared to the Saccharin 1x (1hr) group - 0.5976 ± 0.03555ms. One-Way ANOVA, P=0.0139.

E) Action potential threshold was not significantly different between the groups. Cage control (− 31.83± 2.971 mV), Water (−29.27 ± 2.060 mV), Saccharin 1x (−30.73 ± 2.385 mV), Quinine 1x (− 29.35 ± 3.071 mV), Saccharin 5x (−30.38 ± 2.493 mV), and Saccharin 1x (4hr) (−34.61 ± 2.174 mV). One-Way ANOVA, P= 0.7652.

F) Input resistance was similar among the different treatment groups. Cage control (118.4 ± 9.771 MΩ), Water (136.5 ± 14.40 MΩ), Saccharin 1x (146.6 ±14.22 MΩ), Quinine 1x (139.2 ± 16.86 MΩ), Saccharin 5x (156.1 ± 22.85 MΩ), and Saccharin 1x (4hr) (154.9 ± 22.41 MΩ). One-Way ANOVA, P= 0.6304.

G) SAG ratio was not significantly different between the groups. Cage control (14.91 ± 2.195), Water (8.751± 2.021), Saccharin 1x (11.67±1.790), Quinine 1x (14.15 ± 2.159), Saccharin 5x (11.92 ± 3.395), and Saccharin 1x (4hr) (14.99 ± 2.770). One-Way ANOVA, P= 0.2232.

H) Membrane time constant was significantly different among the treatment groups. Cage control (14.71±1.944 msec), Water (18.03 ± 2.309 msec), Saccharin 1x (14.27 ± 1.666 msec), Quinine 1x (23.21 ± 2.717 msec), Saccharin 5x (17.11 ± 2.296 msec), and Saccharin 1x (4hr) (26.09 ± 5.331msec). One-Way ANOVA, P= 0.0321.

Data are shown as mean ± SEM. *p<0.05, **p<0.01.

Extended Figure 2-1: Learned aversive taste memory retrieval suppresses the excitability of burst spiking LIV-VI aIC-BLA neurons

We compared the intrinsic properties of BS and RS LIV-VI aIC-BLA neurons following Saccharin 2xs (BS=13, RS=7, cells) and CTA memory retrieval (BS=12, RS=15, cells).

A) Excitability in BS LIV-VI aIC-BLA neurons was significantly reduced in the CTA retrieval group compared to Saccharin 2x. Two-Way repeated measures ANOVA, Current x Treatment: p<0.0001.

B) Input resistance in BS LIV-VI aIC-BLA neurons was significantly enhanced in the Saccharin 2x (180.3 ± 15.15 MΩ) compared to CTA retrieval (110.9 ± 12.98 MΩ). Unpaired t test, p= 0.0022.

C) Action potential amplitude in BS LIV-VI aIC-BLA neurons was significantly increased in the CTA retrieval group compared to Saccharin 2x (46.18 ± 1.666 mV), and CTA retrieval (57.87 ± 1.678 mV). Mann Whitney test, p<0.0001.

D) SAG ratio in BS LIV-VI aIC-BLA neurons was significantly suppressed in the Saccharin 2x (7.017 ± 1.317) compared to CTA retrieval (16.8 ± 1.869). Mann Whitney test, p= 0.0005.

E) Representative traces of RS LIV-VI aIC-BLA neurons firing from the two treatments. Scale bars 20 mV vertical and 50ms horizontal in response to 150 pA step current.

F) Excitability in RS LIV-VI aIC-BLA neurons was similar in the CTA retrieval and Saccharin 2x. Two-Way repeated measures ANOVA, Current x Treatment: p= 0.0953.

G) Input resistance in RS LIV-VI aIC-BLA neurons was not significantly different in between the groups. Saccharin 2x (182.6 ± 19.62 MΩ), and CTA retrieval (156.7 ± 10.11 MΩ). Mann Whitney test, p >0.9999.

H) SAG ratio in RS LIV-VI aIC-BLA neurons was not significantly different between the groups. Saccharin 2x (9.297 ± 2.347), and CTA retrieval (10.71 ± 1.536). Mann Whitney test, p= 0.5815.

I) Action potential amplitude in RS LIV-VI aIC-BLA neurons was not significantly different between the groups. Saccharin 2x (54.62 ± 2.058 mV), and CTA retrieval (54.89 ± 1.13 mV). Mann Whitney test, p>0.9999.

J) AP half-width in RS LIV-VI aIC-BLA neurons was significantly reduced following CTA memory retrieval (0.5633 ± 0.01703msec) compared to the Saccharin 2x (0.6614 ± 0.04149msec). Mann Whitney test, p= 0.0200.

K) Membrane time constant was similar in both treatment groups. Saccharin 2x RS (18.36 ± 2.842ms), and CTA memory retrieval RS (24.08 ± 2.023msec). Mann Whitney test, p= 0.0777.

Data are shown as mean ± SEM. *p<0.05, **p<0.01, ***p<0.001, ****p<0.0001.

Extended Figure 3-1: Extinction of CTA enhances, excitability of burst spiking LIV-VI aIC-BLA projecting neurons

We compared the intrinsic properties of BS and RS LIV-VI aIC-BLA neurons following the Extinction (BS=11, RS=3, cells) and Reinstatement (BS=10, RS=5, cells).

A) Excitability in BS LIV-VI aIC-BLA was significantly enhanced in Extinction group comparing to Reinstatement. Two-Way repeated measures ANOVA, Current x Treatment: p<0.0001.

B) sAHP in BS LIV-VI aIC-BLA neurons was significantly enhanced in the Extinction group (−2.104 ± 0.4466 mV) compared to Reinstatement (−3.804 ± 1.339 mV) neurons. Mann Whitney test, p=0.0230.

C) Action potential threshold in BS LIV-VI aIC-BLA neurons was significantly reduced in the Extinction group (−37.41 ± 1.636 mV) compared to Reinstatement (−27.5 ± 2.195 mV). Unpaired t test, p= 0.0016.

D) Input resistance in BS LIV-VI aIC-BLA neurons was similar in the two treatment groups. Extinction (131.1 ± 13.93 MΩ) and Reinstatement BS (157.4 ± 10.56 MΩ). Mann Whitney test, p= 0.1321.

E) SAG ratio in BS LIV-VI aIC-BLA neurons was enhanced following Extinction (13.69 ± 1.541) neurons compared to Reinstatement BS (9.124 ± 1.03). Unpaired t test, p= 0.0262.

F) Membrane time constant in BS LIV-VI aIC-BLA neurons was significantly reduced in the Extinction group (14.52 ± 2.714 msec) compared to Reinstatement (26.93 ± 1.893) neurons. Mann Whitney test, p= 0.0062.

G) Representative traces of RS LIV-VI aIC-BLA firing from two treatment groups. Scale bars 20 mV vertical and 50 msec horizontal in response to 150 pA current step.

H) Excitability of RS LIV-VI aIC-BLA neurons in both treatment groups.

I) Input resistance in RS LIV-VI aIC-BLA neurons was similar in the Extinction (224.2 ± 21.29 MΩ) and Reinstatement (221.2 ± 18.9 MΩ) groups.

J) SAG ratio in RS LIV-VI aIC-BLA neurons was not different between the Extinction (7.515 ± 2.666) and Reinstatement (9.486 ± 1.846) groups.

K) Membrane time constant in RS LIV-VI aIC-BLA neurons was not different between the Extinction (28.69 ± 2.138msec) and Reinstatement groups (22.58 ± 2.632msec).

Extended Figure 5-1: PCA showing Burst vs Regular spiking LIV-VI aIC-BLA neurons all range of excitability vs 350 pA only

A) PCA of BS and RS LIV-VI aIC-BLA neurons all range of excitability (50-350 pA and all other intrinsic properties measured). Sampled population across six treatment groups (Saccharin 1x, Saccharin 2x, Saccharin 5x, CTA retrieval, Extinction, Reinstatement).

B) PCA of BS and RS LIV-VI aIC-BLA neurons excitability of 350 pA only and all other intrinsic properties measured. Sampled population across six treatment groups (Saccharin 1x, Saccharin 2x, Saccharin 5x, CTA retrieval, Extinction, Reinstatement).

Extended Figure 5-2: PCA variable contributions and component loadings of BS and RS LIV-VI aIC-BLA projecting neurons

A) Column chart demonstrating the individual and cumulative proportion of the variance accounted by principal components following PCA of BS LIV-VI aIC-BLA projecting neurons in the two groups of treatments (Saccharin 1x, Saccharin 2x, Extinction vs. CTA retrieval, 5x Saccharin, Reinstatement).

B) Table summarizing the contribution of individual variables (loadings) to the coordinate value of the principal components segregating the two groups (score).

C) Communalities table, demonstrating the amount of variance in each variable that is accounted for by the extraction of principal components. Initial communalities are estimates of the variance in each variable accounted for by all components or factors (=1.00).

