## Supplementary figures for "Intrinsic excitability in layer IV-VI anterior insula to basolateral amygdala projection neurons encodes the confidence of taste valence"

A.

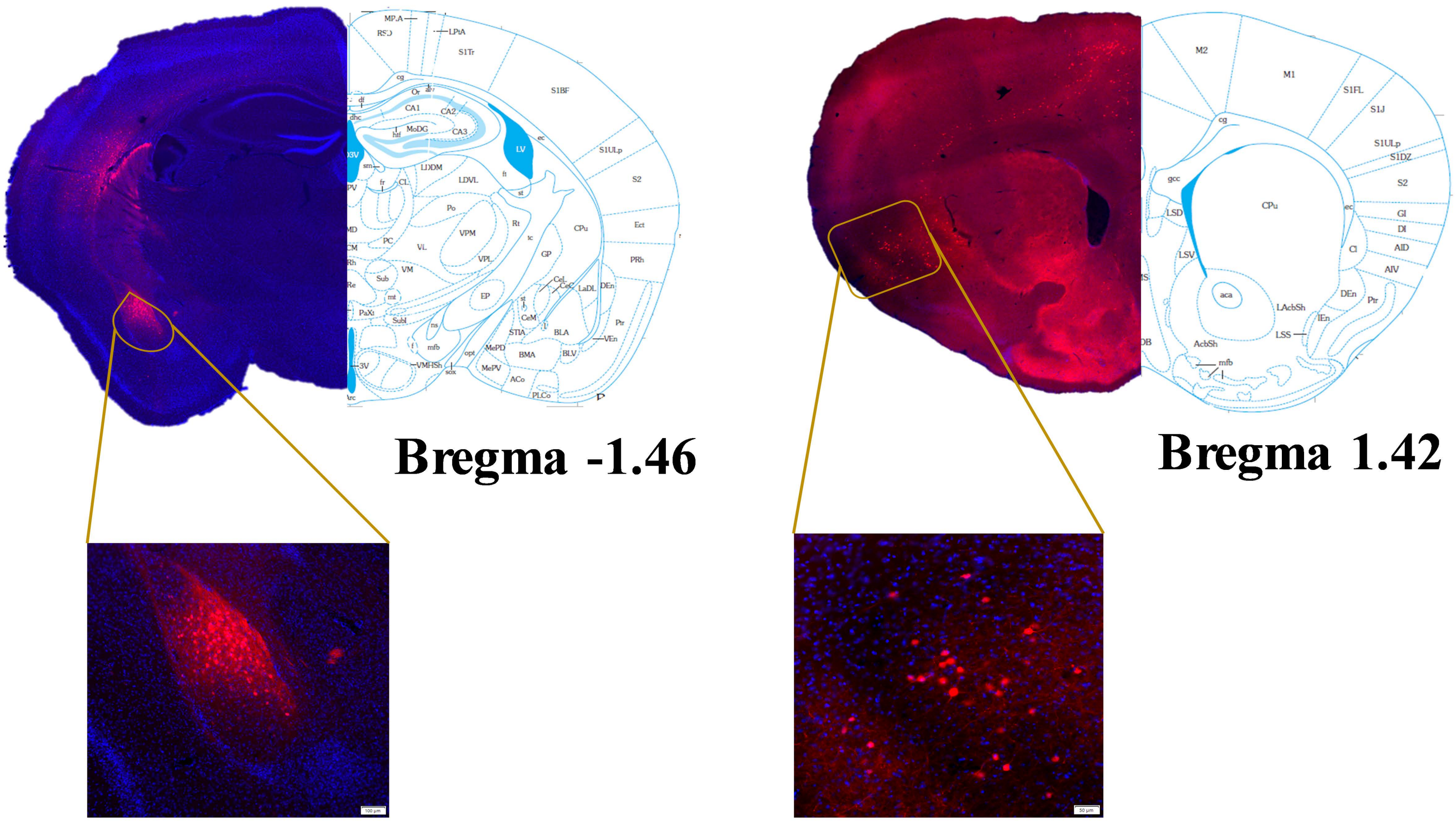


B.

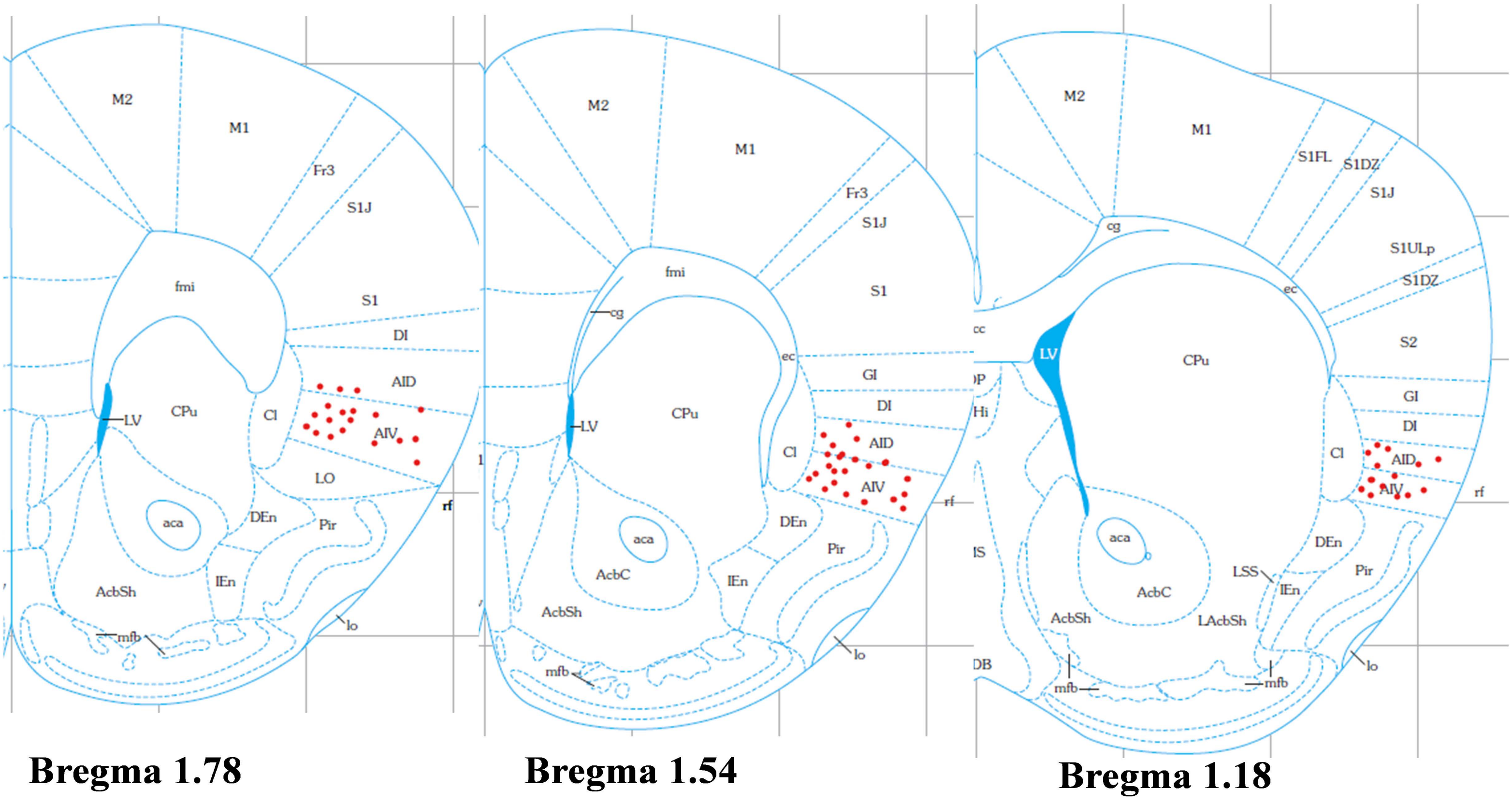

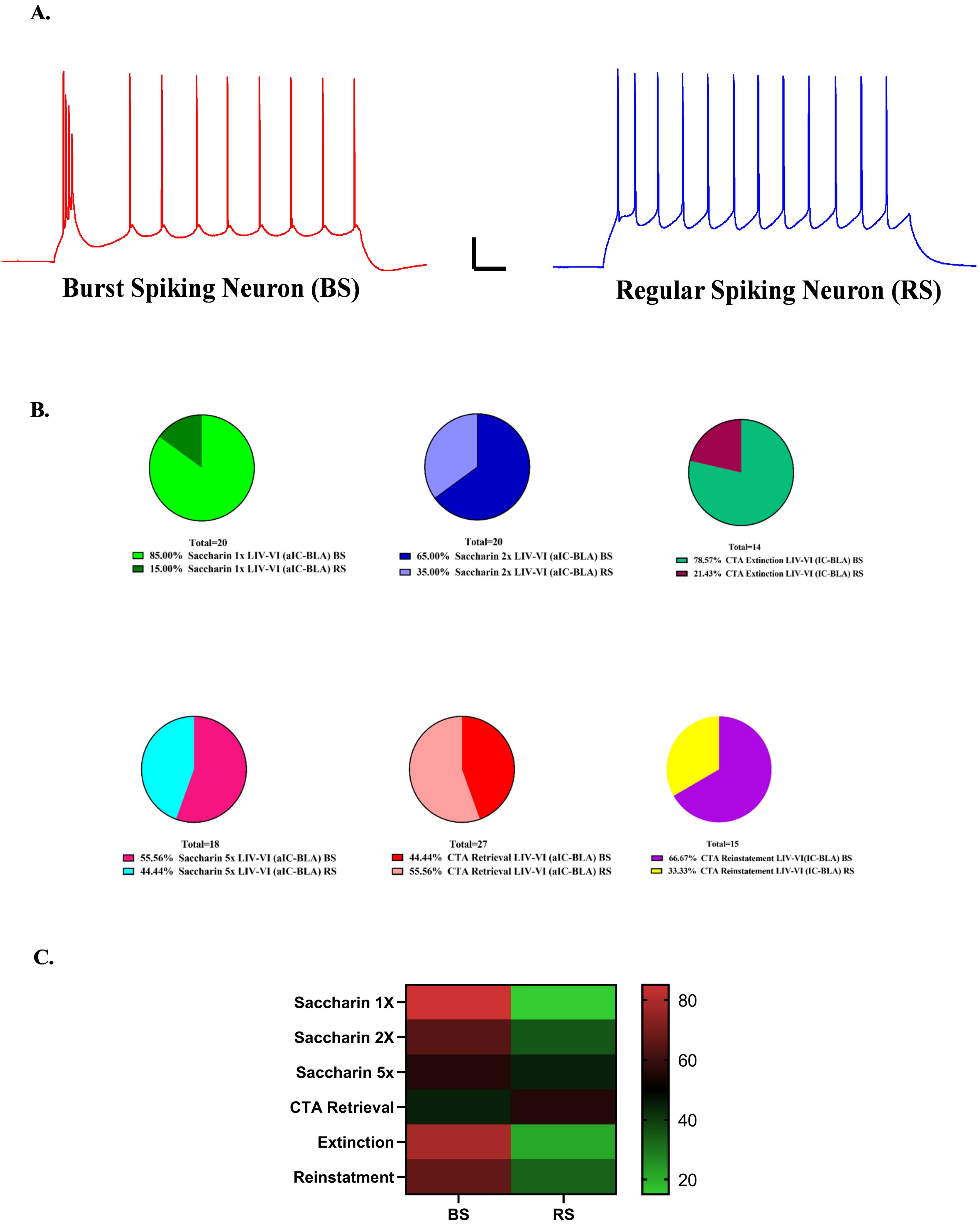
A.
Burst Spiking Neuron (BS)
Regular Spiking Neuron (RS)
B.
Total=20
85.00% Saccharin 1x LIV-VI (aIC-BLA) BS
15.00% Saccharin 1x LIV-VI (aIC-BLA) RS
Total=20
65.00% Saccharin 2x LIV-VI (aIC-BLA) BS
35.00% Saccharin 2x LIV-VI (aIC-BLA) RS
Total=14
78.57% CTA Extinction LIV-VI (IC-BLA) BS
21.43% CTA Extinction LIV-VI (IC-BLA) RS
Total=18
55.56% Saccharin 5x LIV-VI (aIC-BLA) BS
44.44% Saccharin 5x LIV-VI (aIC-BLA) RS
Total=27
44.44% CTA Retrieval LIV-VI (aIC-BLA) BS
55.56% CTA Retrieval LIV-VI (aIC-BLA) RS
Total=15
66.67% CTA Reinstatement LIV-VI(IC-BLA) BS
33.33% CTA Reinstatement LIV-VI (IC-BLA) RS
C.
Saccharin 1X
Saccharin 2X
Saccharin 5x
CTA Retrieval
Extinction
Reinstatment
BS
RS
80
60
40
20

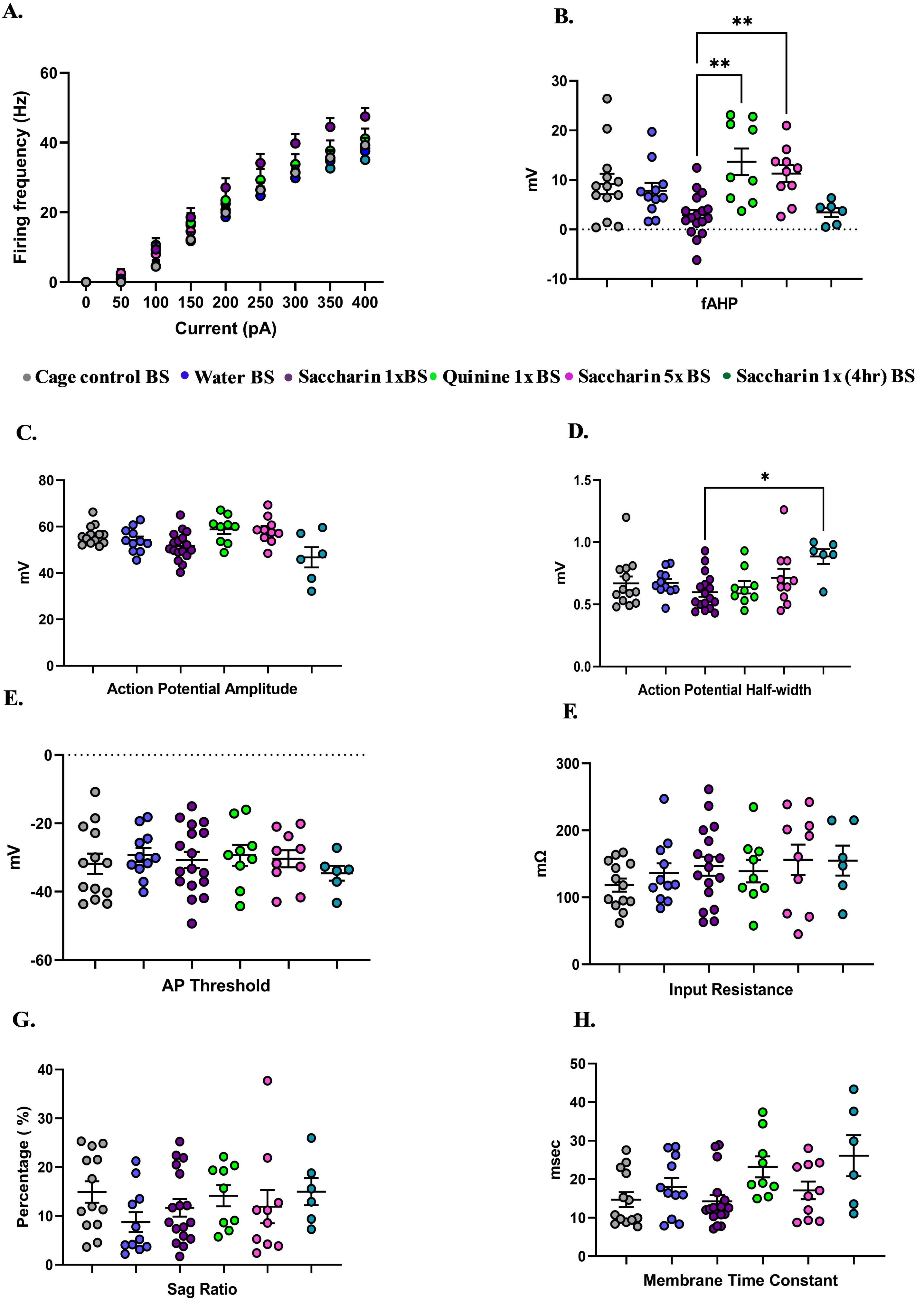
A.
B.
C.
D.
E.
F.
G.
H.
Firing frequency (Hz)
Current (pA)
0
50
100
150
200
250
300
350
400
20
40
60
mV
fAHP
**
**
-10
10
30
Cage control BS
Water BS
Saccharin 1xBS
Quinine 1x BS
Saccharin 5x BS
Saccharin 1x (4hr) BS
Action Potential Amplitude
80
Action Potential Half-width
*
0.0
0.5
1.0
1.5
AP Threshold
-20
-40
-60
Input Resistance
mΩ
300
Sag Ratio
Percentage ( %)
Membrane Time Constant
msec
50

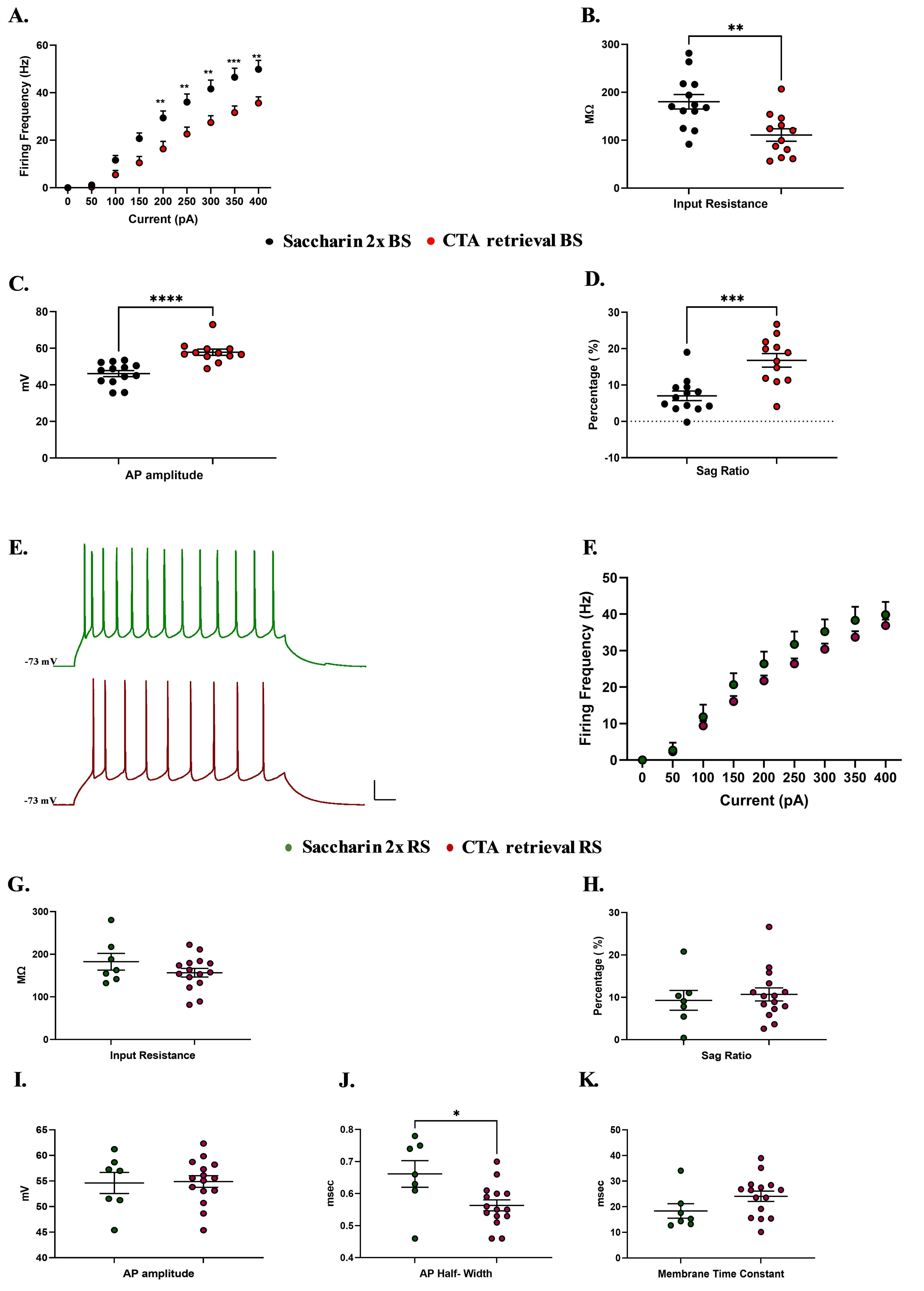

A.
Firing Frequency (Hz)
Current (pA)
B.
Input Resistance
MΩ
Saccharin 2x BS
CTA retrieval BS
C.
AP amplitude
mV
D.
Sag Ratio
Percentage ( %)
E.
-73 mV
-73 mV
F.
Firing Frequency (Hz)
Current (pA)
Saccharin 2x RS
CTA retrieval RS
G.
Input Resistance
MΩ
H.
Sag Ratio
Percentage ( %)
I.
AP amplitude
mV
J.
AP Half- Width
msec
K.
Membrane Time Constant
msec

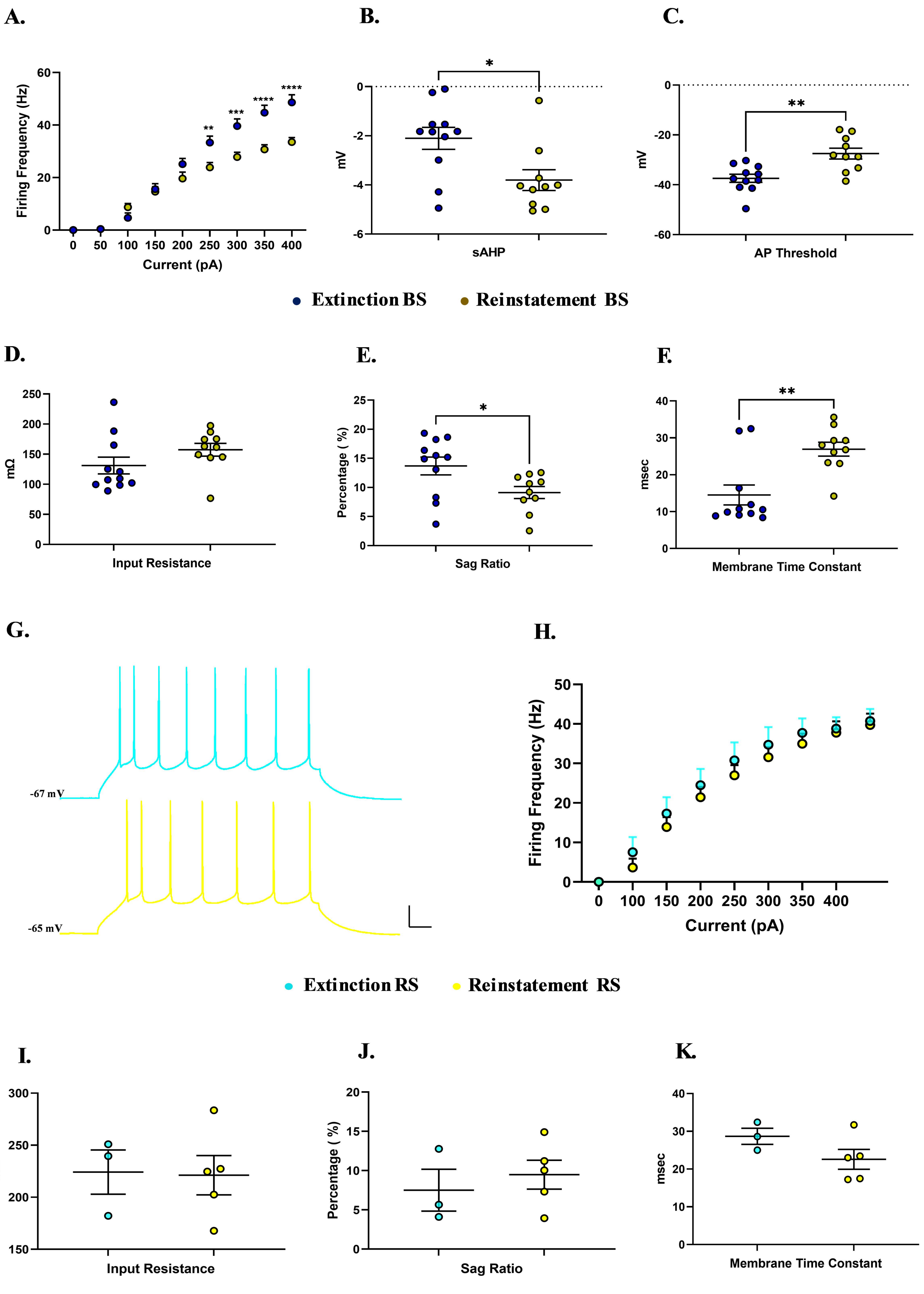
A.
B.
C.
Firing Frequency (Hz)
Current (pA)
0
50
100
150
200
250
300
350
400
**
***
****
****
mV
sAHP
*
AP Threshold
**
Extinction BS
Reinstatement BS
D.
E.
F.
mΩ
Input Resistance
Percentage ( %)
Sag Ratio
*
msec
Membrane Time Constant
**
G.
H.
-67 mV
-65 mV
Firing Frequency (Hz)
Current (pA)
Extinction RS
Reinstatement RS
I.
J.
K.
mΩ
Input Resistance
Percentage ( %)
Sag Ratio
msec
Membrane Time Constant

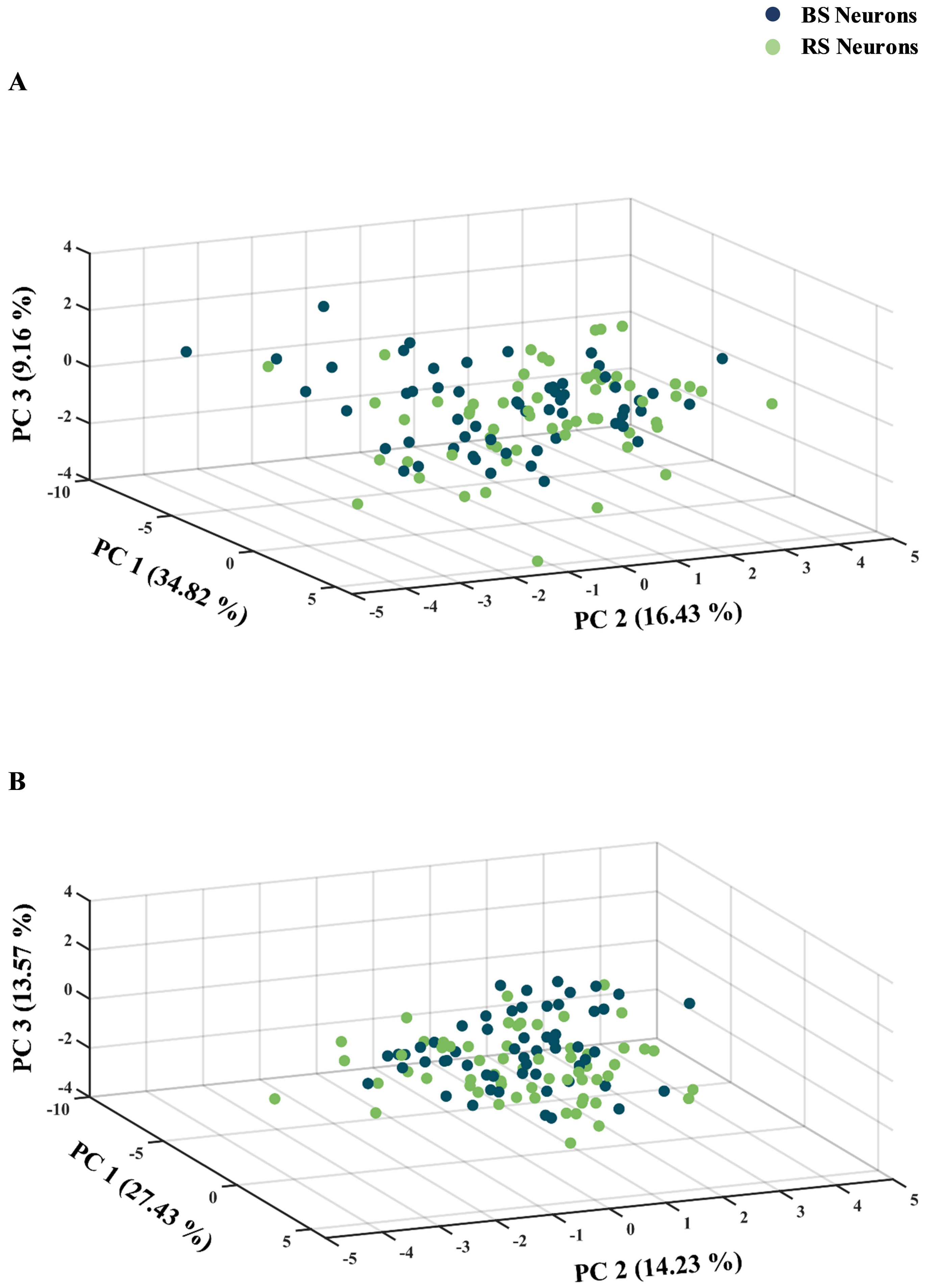
BS Neurons
RS Neurons
A
PC 3 (9.16 %)
PC 1 (34.82 %)
PC 2 (16.43 %)
B
PC 3 (13.57 %)
PC 1 (27.43 %)
PC 2 (14.23 %)

A

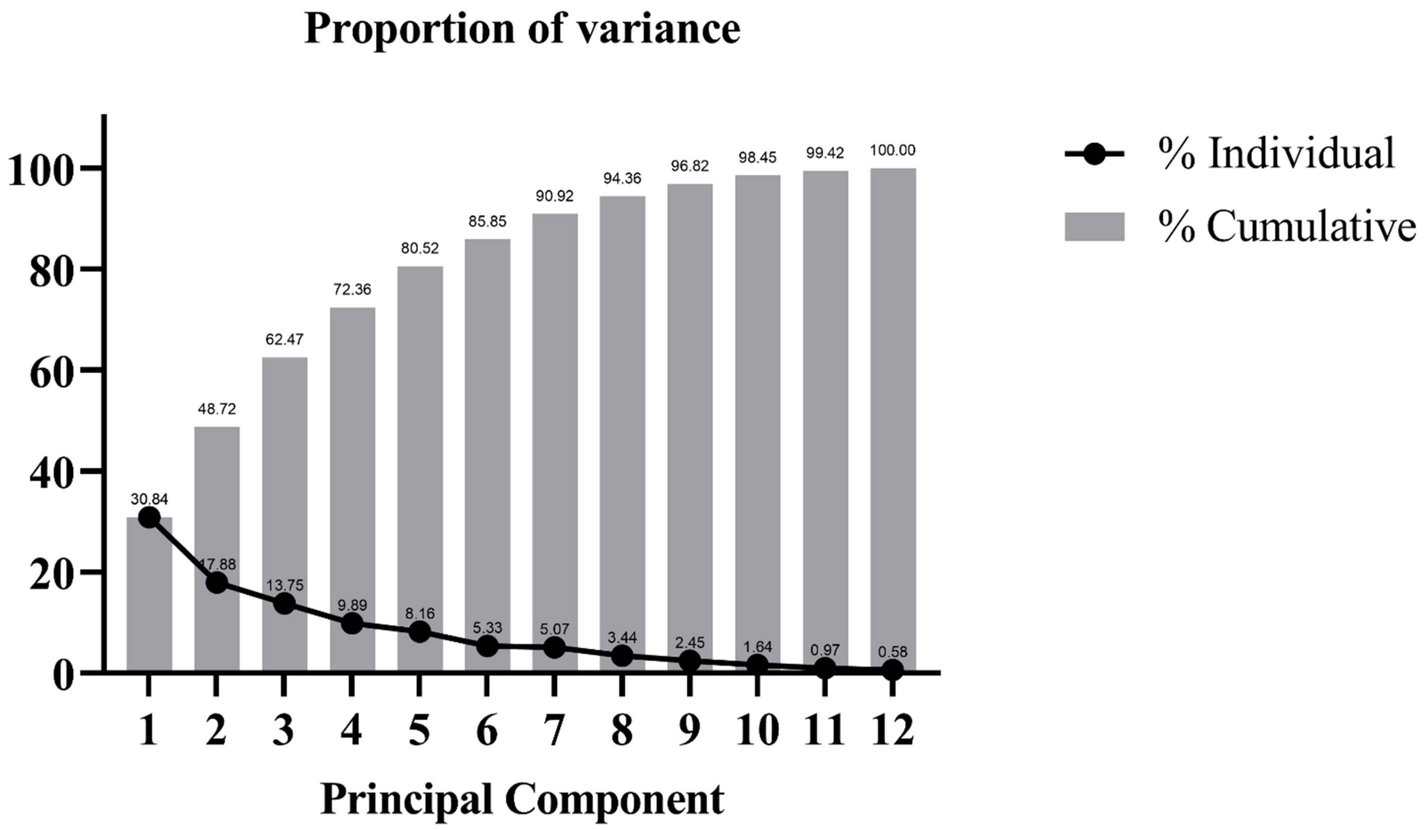


B

**Principal component analysis (PCA) loadings**

| Variables | PC1 | PC2 | PC3 |
|---|---|---|---|
| **350pA** | 0.503981 | **-0.69587** | -0.22966 |
| **RMP(mV)** | 0.221503 | -0.03489 | **0.718035** |
| **mAHP (mV)** | **-0.82421** | -0.05264 | **-0.6095** |
| **sAHP(mV)** | **-0.85985** | -0.17996 | -0.09883 |
| **fAHP(mV)** | 0.043537 | **0.682681** | -0.13433 |
| **IR(MΩ)** | **0.694764** | 0.078472 | -0.45433 |
| **Sag Ratio** | -0.09223 | -0.18107 | **0.889949** |
| **Time constants(ms)** | 0.157048 | 0.477532 | -0.35429 |
| **AP amplitude (mV)** | -0.16953 | 0.493821 | 0.350324 |
| **AP Halfwidth(ms)** | 0.132945 | **0.614103** | -0.0942 |
| **AP threshold(mV)** | 0.274889 | 0.499891 | -0.16142 |
| **Rheobase(pA)** | **-0.88304** | 0.089672 | 0.043254 |

C

**Communalities**

| Variables | Initial | Extraction |
|---|---|---|
| **Exc350p** | 1.000 | 0.859 |
| **RMP** | 1.000 | 0.496 |
| **mAHP** | 1.000 | 0.856 |
| **sAHP** | 1.000 | 0.860 |
| **fAHP** | 1.000 | 0.503 |
| **IR** | 1.000 | 0.897 |
| **Sag_Ratio** | 1.000 | 0.867 |
| **Time_constant** | 1.000 | 0.732 |
| **AP_amplitude** | 1.000 | 0.650 |
| **AP_Halfwidth** | 1.000 | 0.741 |
| **AP_Threshold** | 1.000 | 0.426 |
| **Rheobase** | 1.000 | 0.795 |

| Groups | RMP (mV) | fAHP (mV) | mAHP (mV) | sAHP (mV) | Input Resistance (MΩ) | Sag ratio (%) | Time constant (msec) | AP threshold (mV) | AP Amplitude (mV) | AP half-width (msec) | Rheobase (pA) |
|---|---|---|---|---|---|---|---|---|---|---|---|
| **LIV-VI aIC-BLA Cage control** | -68.28 ± 0.8506 (19) | 9.191 ± 1.449 (19) ** ## | -3.73 ± 0.4241 (19) | -1.881 ± 0.3376 (19) | 120.7 ± 7.686 (19) | 12.41 ± 1.938 (19) | 15.03 ± 1.376 (19) | -30.76 ± 2.139 (19) | 56.28 ± 0.8818 (19) ^^ | 0.6774 ± 0.03816 (19) | 87.47 ± 9.127 (19) |
| **LIV-VI aIC-BLA Water** | -69.3 ± 0.9051 (23) | 8.15 ± 0.8288 (23) ** | -5.535 ± 0.6754 (23) | -3.277 ± 0.4603 (23) | 139.1 ± 9.021(23) | 8.909 ± 1.306 (23) | 19.24 ± 1.62 (23) | -31.05 ± 1.30 (23) | 52.43 ± 1.034 (23) | 0.6243 ± 0.02021 (23) | 85 ± 11.76 (23) |
| **LIV-VI aIC-BLA Saccharin 1x** | -69.21 ± 0.94 (20) | 3.016 ± 0.9423 (20) #### | -4.17 ± 0.4542 (20) | -2.521 ± 0.2735 (20) | 145.8 ± 12.56 (20) | 10.89 ± 1.621 (20) | 14.82 ± 1.485 (20) # | -30.9 ± 2.141 (20) | 52.03 ± 1.308 (20) | 0.6005 ± 0.0326 (20) | 74.25 ± 11.39 (20) |
| **LIV-VI aIC-BLA Quinine 1x** | -67.32 ± 1.092 (19) | 13.56± 1.562 (19) $$$$ ~ | -5.858 ± 0.5613 (18) | -3.634 ± 0.3632 (18) | 146 ± 9.094 (19) | 12.13 ± 1.23 (19) | 21.55 ± 1.638 (19) | -29.74 ± 1.989 (19) | 57.11 ± 1.376 (19) *^^^ | 0.6 ± 0.03555 (19) | 69.89 ± 8.932 (19) |
| **LIV-VI aIC-BLA Saccharin 5x** | -66.53 ± 1.358 (18) | 8.158 ± 1.356 (18) ** ### | -3.999 ± 0.653 (18) | -2.695 ± 0.5083 (18) | 144.6 ± 14.68 (18) | 9.392 ± 2.127 (18) | 17.3 ± 1.66 (18) | -32.66 ± 1.783 (18) | 56.48 ± 1.337 (18) | 0.6539 ± 0.04814 (18) | 78.61 ± 10.75 (18) |
| **LIV-VI aIC-BLA Saccharin 1x (4hr)** | -69.47 ± 0.7569 (17) | 5.989 ± 1.074 (17) ** | -4.411 ± 0.8962 (17) | -2.631 ± 0.6949 (17) | 153.9 ± 11.10 (17) | 10.97 ± 1.475 (17) | 26.21 ± 2.421 (17) ** $ $ | -31.03 ± 1.511 (17) | 52.24 ± 2.311 (17) | 0.7765 ± 0.03641 (17) ** | 78.88 ± 9.274 (17) |
| **LIV-VI aIC-BLA Saccharin 2x** | -70.79 ± 1.242 (20) | 5.223 ± 0.8217 (20) ### | -5.301 ± 0.7863 (20) | -3.351 ±0.3798 (20) | 181.1 ± 11.7 (20) | 7.815 ± 1.176 (20) | 19.28 ± 1.837 (20) | -32.85 ± 1.447 (20) | 49.14 ± 1.568 (20) ### | 0.578 ± 0.02994 (20) | 69.6 ± 10.71 (20) |
| **LIV-VI aIC-BLA CTA Retrieval** | -68.78 ± 0.8419 (27) | 7.97 ± 1.018 (27) ## | -5.213 0.4544 (27) | -2.69 ± 0.3064 (27) | 136.4 ± 9.064 (27) ^^ | 13.41 ± 1.31 (27) ^^ | 20.96 ± 1.724 (27) * | -31.61 ± 2.68 (27) | 56.21 ± 0.9978 (27) ^^^ | 0.5959 ± 0.0208 (27) | 90.44 ± 17.56 (27) |
| **LIV-VI aIC-BLA Extinction** | -65.98 ± 1.457 (14) | 4.731 ± 1.021 (14) | -5.076 ± 0.6981 (14) | -2.895 ± 0.6547 (14) | 151.1 ± 15.63 (14) | 12.37 ± 1.471 (14) | 17.55 ± 2.684 (14) ~ | -36.06 ± 1.481 (14) ~ | 55.09 ± 2.122 (14) ^ | 0.7386 ± 0.03145 (14) | 69.21 ± 7.454 (14) |
| **LIV-VI aIC-BLA Reinstatement** | -68.57 ± 0.936 (15) | 5.932 ± 1.292 (15) | -5.673 ± 0.4288 (15) | -3.612 ±0.3033 (15) | 178.7 ± 12.1 (15) | 9.245 ± 0.884 (15) | 25.48 ± 1.58 (15) ** | -29.43 ± 1.731 (15) | 53.1 ± 2.906 (15) | 0.8187 ± 0.06929 (15) | 67.6 ± 8.753 (15) |

* vs. Saccharin 1X

^ vs. Saccharin 2x

$ vs. Cage Control

### vs. Quinine

~ vs. Reinstatement

#### Table 1: Summary of LIV-VI aIC-BLA intrinsic properties

Values are expressed in mean ± SEM. The number of cells is in parentheses. Statistical analysis was performed by One-way ANOVA Post-hoc Tukey's and Dunn's multiple comparisons. Student's t-test was performed for the comparison between two groups. RMP - resting membrane potential, fAHP, mAHP and sAHP - fast, medium, slow after hyperpolarization potentials, respectively. AP Thresh - action potential threshold, AP Amp - action potential amplitude. AP half-width - action potential half-width.

| Groups | RMP (mV) | mAHP (mV) | Input resistance (MΩ) | Sag ratio (%) | Time constant (msec) | AP thresh (mV) | AP Amp (mV) | AP half-width (msec) | Rheobase (pA) |
|---|---|---|---|---|---|---|---|---|---|
| **LI-III aIC-BLA Water** | -70.84 ± 1.09 (14) | -2.411 ± 0.6767 (14) | 136.4 ± 16.85 (14) | 3.172 ± 1.082 (14) | 11.17 ± 1.169 (14) | -31.35 ± 1.547 (14) | 56 ± 1.719 (14) | 0.6557 ± 0.03222 (14) | 140.7 ± 31.83 (14) |
| **LI-III aIC-BLA Saccharin 1x** | -72.27 ± 1.212 (15) | -1.752 ± 0.4953 (15) | 131.6 ± 13.14 (15) | 3.997 ± 0.9166 (15) | 11.56 ± 1.672 (15) | -31.44 ± 1.621 (15) | 56.33 ± 1.323 (15) | 0.684 ± 0.02767 (15) | 136.1 ± 26.44(15) |
| **LI-III aIC-BLA Saccharin 2x** | -73.44 ± 1.295 (15) | -2.165 ± 0.684 (15) | 142.5 ± 13.17 (15) | 5.793 ± 1.449 (15) | 18.63 ± 1.9 (15) | -31.47 ± 2.511 (15) | 49.76 ± 2.065 (15) | 0.564 ± 0.03708 (15) | 104.7 ± 25.03 (15) |
| **LI-III aIC-BLA CTA Retrieval** | -72 ± 1.117 (17) | -3.087 ± 2.914 (17) | 171.1 ± 22.28 (17) | 6.307 ± 1.368 (17) | 18.94 ± 1.667 (17) | -34.24 ± 1.445 (17) | 48.58 ± 1.472 (17) | 0.5518 ± 0.02698 (17) | 114.3 ± 31.57 (17) |

#### Table 2. Summary of LI-III aIC-BLA intrinsic properties

Values are expressed as mean ± SEM. The number of cells is in parentheses. Statistical analysis was performed by Student's t-test. RMP - resting membrane potential, mAHP - medium after hyperpolarization potentials. AP Thresh - action potential threshold, AP Amp - action potential amplitude. AP half-width - action potential half-width.

| Groups | RMP (mV) | fAHP (mV) | mAHP (mV) | sAHP (mV) | Input resistance (MΩ) | Sag ratio (%) | Time constant (msec) | AP thresh (mV) | AP Amp (mV) | AP half-width (msec) | Rheobase (pA) |
|---|---|---|---|---|---|---|---|---|---|---|---|
| **BS LIV-VI aIC-BLA Cage control** | -68.28 ± 0.9705 (13) | 9.192 ± 2.061 (13) | -4.194 ± 0.5072 (13) | -2.075 ± 0.4500 (13) | 118.4 ± 9.771 (13) | 14.91 ± 2.195 (13) | 14.71±1.944 (13) | -31.83± 2.971 (13) | 56.27 ±1.147 (13) | 0.6692 ± 0.05460 (13) | 74.54 ± 8.471 (13) |
| **BS LIV-VI aIC-BLA Water** | -69.00 ± 1.639 (11) | 7.800 ± 1.607(11) | -4.870 ± 0.8838 (11) | -2.826 ± 0.6069 (11) | 136.5 ± 14.40 (11) | 8.751± 2.021 (11) | 18.03 ± 2.309 (11) | -29.27 ± 2.060 (11) | 54.21 ±1.572 (11) | 0.6736 ± 0.03111 (11) | 85.91 ±13.44 (11) |
| **BS LIV-VI aIC-BLA Saccharin 1x** | -68.80 ± 1.065 (17) | 2.870 ±1.044 (17) | -4.339 ±0.5083 (17) | -2.564 ±0.3032 (17) | 146.6 ±14.22 (17) | 11.67±1.790 (17) | 14.27 ± 1.666 (17) | -30.73 ± 2.385 (17) | 51.64 ± 1.473 (17) | 0.5976 ± 0.03555 (17) | 63.65 ± 7.679 (17) |
| **BS LIV-VI aIC-BLA Quinine 1x** | -67.37 ± 1.682 (9) | 13.67 ±2.681(9) ## | -6.131 ± 0.6514 (8) | -3.58 ± 0.5788 (8) | 139.2 ± 16.86 (9) | 14.15 ± 2.159 (9) | 23.21 ± 2.717 (9) | -29.35 ± 3.071 (9) | 58.86 ± 2.003 (9) | 0.6378 ± 0.0491 (9) | 59.56 ± 12.28 (9) |
| **BS LIV-VI aIC-BLA Saccharin 5x** | -67.20 ±1.624 (10) | 11.30 ±1.727 (10) ## | -5.174 ±0.8427 (10) | -3.609 ± 0.7205 (10) | 156.1 ± 22.85 (10) | 11.92 ± 3.395 (10) | 17.11 ± 2.296 (10) | -30.38 ± 2.493 (10) | 58.40 ± 1.812 (10) | 0.7140 ± 0.07349 (10) | 76.60 ± 15.32 (10) |
| **BS LIV-VI aIC-BLA Saccharin 1x (4hr)** | -68.20 ±1.293 (6) | 3.433 ± 0.9245(6) | -6.693 ±1.442 (6) | -3.130 ± 1.637 (6) | 154.9 ± 22.41(6) | 14.99 ± 2.770 (6) | 26.09 ± 5.331 (6) | -34.61 ± 2.174 (6) | 46.79 ± 4.359 (6) | 0.8850 ± 0.05943 (6) # | 60.83 ± 11.36 (6) |
| **BS LIV-VI aIC-BLA Saccharin 2x** | -71.33 ± 1.641 (13) | 4.169 ± 0.9225 (13) | -5.368 ± 0.9616 (13) | -3.226 ± 0.4899 (13) | 180.3 ± 15.15 (13) ** | 7.017 ± 1.317 (13) *** | 19.77 ± 2.447 (13) | -34.20 ± 1.987 (13) | 46.18 ± 1.666 (13) *** | 0.5331 ± 0.03522 (13) | 77.54 ± 15.69 (13) |
| **BS LIV-VI aIC-BLA CTA Retrieval** | -67.37 ± 1.21 (12) | 5.473 ± 1.464 (12) | -4.633 ± 0.5831 (12) | -1.932 ± 0.4462 (12) | 110.9 ± 12.98 (12) | 16.8 ± 1.869 (12) | 17.06 ± 2.608 (12) | -34.28 ± 1.771 (12) | 57.87 ± 1.678 (12) | 0.6367± 0.03961 (12) | 88.75 ± 9.847 (12) |
| **BS LIV-VI aIC-BLA Extinction** | -67.36 ± 1.43 (11) | 3.943 ±1.111 (11) | -4.816 ± 0.8447 (11) | -2.104 ± 0.4466 (11) ~ | 131.1 ± 13.93 (11) | 13.69 ± 1.541 (11) | 14.52 ± 2.714 (11) ~ ~ | -37.41 ± 1.636 (11) ~ ~ | 57.3 ± 2.023 (11) | 0.7155 ± 0.03674 (11) | 81 ± 6.932 (11) |
| **BS LIV-VI aIC-BLA Reinstatement** | -68.88 ± 1.163 (10) | 6.432 ± 1.737 (10) | -5.243 ± 0.5853 (10) | -3.804 ± 1.339 (10) | 157.4 ± 10.56 (10) | 9.124 ± 1.03 (10) | 26.93 ± 1.893 (10) | -27.5 ± 2.195 (10) | 57.36 ± 3.001 (10) | 0.769 ± 0.03494 (10) | 73.9 ± 12.45 (10) |

*** BS LV/VI aIC-BLA Saccharin 2x vs. CTA Retrieval, *p<0.05, ** p<0.01, *** p<0.001.**

**~ BS LV/VI aIC-BLA Extinction vs. Reinstatement, ~ p<0.05, ~ ~ p<0.01, ~ ~ ~ p<0.001.**

**# BS LV/VI aIC-BLA Saccharin 1x vs. Quinine 1x, Saccharin 5x and Saccharin 1x (4hr), # p<0.05, ## p<0.01.**

#### Table 3. Summary of BS LIV-VI aIC-BLA intrinsic properties

Values are expressed in mean ± SEM. The number of cells is in parentheses. Statistical analysis was performed by One-way ANOVA Post-hoc Tukey's and Dunn's multiple comparisons. Student's t-test was performed for the comparison between two groups. RMP - resting membrane potential, fAHP, mAHP and sAHP - fast, medium, slow after hyperpolarization potentials, respectively. AP Thresh - action potential threshold, AP Amp - action potential amplitude. AP half-width - action potential half-width.

| Groups | RMP (mV) | fAHP (mV) | mAHP (mV) | sAHP (mV) | Input resistance (MΩ) | Sag ratio (%) | Time constant (msec) | AP thresh (mV) | AP Amp (mV) | AP half-width (msec) | Rheobase (pA) |
|---|---|---|---|---|---|---|---|---|---|---|---|
| **RS LIV-VI aIC-BLA Cage control** | -68.29 ±1.830 (6) | 9.188 ± 1.351 (6) | -2.725 ± 0.6462 (6) | -1.460 ± 0.4409 (6) | 125.7 ± 13.03 (6) | 6.993 ± 3.030 (6) | 15.73 ± 1.333 (6) | -28.44 ± 2.166 (6) | 56.30 ± 1.422 (6) | 0.6950 ± 0.03170 (6) | 115.5 ± 18.62 (6) |
| **RS LIV-VI aIC-BLA Water** | -69.57 ±0.9422 (12) | 8.472 ± 0.6792 (12) | -6.144 ± 1.013 (12) | -3.690 ± 0.6876 (12) | 141.5 ±11.75 (12) | 9.735 ± 1.400 (12) | 20.35 ± 2.321 (12) | -32.67 ± 1.561 (12) | 50.79 ± 1.238 (12) | 0.5792 ± 0.01928 (12) | 84.17 ± 19.48 (12) |
| **RS LIV-VI aIC-BLA Saccharin 1x** | -71.57 ± 1.120 (3) | 3.840 ± 2.530 (3) | -3.210 ± 0.9005 (3) | -2.277 ± 0.7297 (3) | 141.3 ±28.45 (3) | 6.455 ± 3.099 (3) | 17.95 ± 2.856 (3) | -31.86 ± 5.650 (3) | 54.23 ± 2.660 (3) | 0.6167 ± 0.09939 (3) | 134.3 ± 58.52 (3) |
| **RS LIV-VI aIC-BLA Quinine 1x** | -67.28 ±1.505 (10) | 11.63 ± 1.616 (10) | -5.639 ± 0.8918 (10) | -3.678 ± 0.4895 (10) | 151.4 ± 9.055 (10) | 10.31 ± 1.115 (10) | 23.6 ± 1.77(10) | -30.1 ± 2.732 (10) | 55.53 ± 1.845 (10) | 0.6230 ± 0.0535 (10) | 79.20 ± 12.73 (10) |
| **RS LIV-VI aIC-BLA Saccharin 5x** | -65.70 ± 2.378 (8) | 4.235 ± 1.141 (8) | -2.530 ± 0.7960 (8) | -1.553 ±0.4915 (8) | 130.1 ± 16.85 (8) | 6.237 ± 1.910 (8) | 17.54 ± 2.561 (8) | -35.50 ± 2.302 (8) | 54.09 ± 1.734 (8) | 0.5788 ± 0.05034 (8) | 81.13 ±15.88 (8) |
| **RS LIV-VI aIC-BLA Saccharin 1x (4hr)** | -70.17 ± 0.9075 (11) | 7.383 ± 1.439 (11) | -3.165 ± 0.9897 (11) | -2.359 ± 0.6647 (11) | 153.3 ± 12.96 (11) | 8.782 ± 1.389 (11) | 26.28 ± 2.596 (11) | -28.43 ± 1.652 (11) | 51.66 ± 3.053 (11) | 0.7773 ± 0.04702 (11) | 88.73 ± 12.25 (11) |
| **RS LIV-VI aIC-BLA Saccharin 2x** | -69.79 ± 1.921 (7) | 7.180 ± 1.402 (7) | -5.016 ± 1.460 (7) | -3.581 ± 0.6324 (7) | 182.6 ± 19.62 (7) | 9.297 ± 2.347 (7) | 18.36 ± 2.842 (7) | -30.36 ± 1.638 (7) | 54.62 ± 2.058 (7) | 0.6614 ± 0.04149 (7) | 54.86 ± 8.207 (7) |
| **RS LIV-VI aIC-BLA CTA Retrieval** | -69.9 ± 1.116 (15) | 9.967 ± 1.216 (15) | -5.667 ± 0.6647 (15) | -3.297 ± 0.3599(15) | 156.7 ± 10.11 (15) | 10.71 ± 1.536 (15) | 24.08 ± 2.023 (15) | -33.9 ± 1.132 (15) | 54.89 ± 1.13 (15) | 0.5633 ± 0.01703 (15) * | 91.8 ± 31.15 (15) |
| **RS LIV-VI aIC-BLA Extinction** | -60.93 ± 3.263 (3) | 7.620 ± 1.907 (3) | -6.027 ± 1.062 (3) | -5.797 ± 1.997 (3) | 224.2 ± 21.29 (3) | 7.515 ± 2.666 (3) | 28.69 ± 2.138 (3) | -31.1 ± 1.372 (3) | 46.98 ± 4.432 (3) | 0.8233 ± 0.02603 (3) | 36.67 ± 13.33 (3) |

| | | | | | | | | | | | |
|---|---|---|---|---|---|---|---|---|---|---|---|
| **RS LIV-VI aIC-BLA Reinstatement** | -67.95 ± 1.725 (5) | 4.932 ± 1.893 (5) | -6.532 ± 0.3344 (5) | -3.228 ± 0.3214 (5) | 221.2 ± 18.9 (5) | 9.486 ± 1.846 (5) | 22.58 ± 2.632 (5) | -33.31 ± 2.035 (5) | 44.58 ± 4.569 (5) | 0.918 ± 0.203 (5) | 55 ± 6.885 (5) |

*** RS LV/VI aIC-BLA Saccharin 1x vs. CTA Retrieval, *$p<0.05$.**

#### Table 4: Summary of RS LIV-VI aIC-BLA intrinsic properties

Values are expressed in mean ± SEM. The number of cells is in parentheses. Statistical analysis was performed by Student's t-test was performed for the comparison of two groups. RMP - resting membrane potential, fAHP, mAHP and sAHP - fast, medium, slow after hyperpolarization potentials, respectively. AP Thresh - action potential threshold, AP Amp - action potential amplitude. AP half-width - action potential half-width.
